# β-arrestin-dependent ERK signaling positively correlates with reduced anxiety-like and conditioned fear-related behavior in mice

**DOI:** 10.1101/790568

**Authors:** Mee Jung Ko, Terrance Chiang, Arbaaz A. Mukadam, Grace E. Mulia, Anna M. Gutridge, Angel Lin, Julia A. Chester, Richard M. van Rijn

## Abstract

Exposure to anxiety- or fear-invoking stimuli initiates a convergence of executive actions orchestrated by multiple proteins and neurotransmitters across the brain. Dozens of G protein-coupled receptors (GPCRs) have been linked to regulation of fear and anxiety. GPCR signaling involves canonical G protein pathways but may also engage downstream kinases and effectors through β-arrestin scaffolds. Here, we investigate whether β-arrestin signaling can regulate anxiety-like and fear-related behavior. Using the δ-opioid receptor (δOR) as a model GPCR, we found that β-arrestin 2-dependent activation of extracellular signal–regulated kinases (ERK1/2) in the dorsal hippocampus and the amygdala are critical for δOR agonist-induced anxiolytic-like effects. In contrast, G protein-mediated δOR signaling was associated with decreased ERK1/2 activity and increased fear-related behavior. Our results also indicate unique contributions for β-arrestin isoforms in modulation of anxiety-like and fear-related behavior. Overall, our findings highlight the significance of non-canonical β-arrestin signaling in the regulation of emotions.

**One sentence summary:** Using pharmacological and genetic strategies, we reveal the importance of non-canonical β-arrestin-mediated G protein-coupled receptor signaling in anxiety-like behaviors.

## Introduction

Anxiety- and fear-related behaviors are evolutionary adaptive behaviors important for human survival. However, excessive stimulation of the neural circuits that regulate anxiety- and fear-related behaviors can manifest as psychiatric disorders, such as post-traumatic stress disorder and phobias. The expression of anxiety- and fear-related behaviors is regulated by complex integration of both internal and external physiological and sensory cues that influence reflexive behavior, cognitive control, and executive functions. Accordingly, behavioral correlates of anxiety and fear are regulated by a harmonious activity of neurotransmitters and cellular actions across many overlapping and distinct neural circuits (*1*).

With regard to cellular actions underlying (patho-)physiological behavior, G protein-coupled receptors (GPCRs) play an important role in neuronal signaling. GPCRs bind neurotransmitters and initiate intracellular signal transduction pathways which ultimately affect neuronal excitability, neurotransmitter release and synaptic plasticity. GPCRs, including serotonergic (*2*), dopaminergic (*3*), adrenergic (*4*), opioidergic (*5*) and corticotropin-releasing factor receptors (*6*) have well-documented roles in the modulation of anxiety and fear and thus are exciting drug targets to treat anxiety disorders.

Although drug development at GPCRs has traditionally focused on the canonical G protein pathways, the prior two decades have introduced arrestin-dependent signaling as a new concept of GPCR signal transduction. In particular, the ‘non-visual’ arrestins 2 and 3, referred here as β-arrestin 1 and 2, respectively, have been studied for their ability to drive G-protein-independent signaling (*7*). The primary role of β-arrestins is to desensitize GPCRs, for example increases β-arrestin 2 expression in mice amygdala, caused by HIV1-tat infection, leads to significant reduction in morphine efficacy in this region (*8*). Beyond desensitization, β-arrestin 2 may also partake in receptor signaling by scaffolding with various kinases. For example, β-arrestin 2 p38 MAP kinase signaling has been linked to the aversive effects of κ-opioid receptor (κOR) agonists (*5, 9*), whereas a β-arrestin 2-GSK3β/AKT scaffold appears to be driving the antipsychotic effects of dopamine D_2_ receptors agonists (*10, 11*). Currently, few studies have investigated how signaling scaffolds involving the β-arrestin isoforms may influence anxiety- and fear-like behavior. It is important to begin to address this gap in our current knowledge of the GPCR modulation of psychiatric behavior, especially since the majority of medications that target GPCRs were developed without consideration of the potential adverse or therapeutic effects of β-arrestin signaling. Yet, it is now possible to develop molecules that preferentially activate or avoid β-arrestin signaling and thus have the potential to treat psychiatric disorders more effectively and with a wider therapeutic window (*12, 13*).

Here, we describe our efforts to elucidate the roles of β-arrestin isoforms in mediating GPCR signaling in relation to the modulation of anxiety and fear-like behavior. We chose to utilize the δ-opioid receptor (δOR) as a model GPCR for multiple reasons. Previous studies have shown that the δOR selective agonist SNC80 is an efficacious recruiter of β-arrestin 1 and 2 proteins (*14-16*), and has anxiolytic-like (*17, 18*) and fear-reducing effects (*17, 19*). Moreover, the δOR-selective agonist TAN67, which is a poor β-arrestin 2 recruiter does not reduce anxiety-like behavior in naïve mice (*20*), providing support for our hypothesis that β-arrestin 2 signaling is correlated with and reduced anxiety-like behavior.

Mitogen activated protein kinases (MAPKs) have been implicated with mood disorders and can scaffold with β-arrestin (*21, 22*). Studies have suggested that MAPK signaling, specifically ERK1/2, in the hippocampus and the basolateral amygdala is required for the acquisition and extinction of fear memory (*23, 24*). Therefore, we further hypothesize that β-arrestin-dependent MAPK signaling may contribute to anxiety-like and fear-related behavior. To test our hypotheses, we assessed the degree to which β-arrestin isoforms and MAPK activation were involved in δOR agonist-mediated modulation of unconditioned anxiety-related behavior and cued-induced fear-related behavior. Our results suggest that ERK1/2 activity is differentially modulated by G protein and β-arrestin signaling and is correlated with anxiety-like and fear-related responses in C57BL/6 mice. We noted that different β-arrestin isoforms were involved in the activation of ERK1/2 across various brain regions, including the striatum, hippocampus and amygdala.

## Results

### Involvement of β-arrestin 2 in the modulation of anxiety-like behavior

In 2016, Astra Zeneca revealed that their novel δOR-selective agonist AZD2327 (**Fig. 1a**) was capable of reducing anxiety-like behavior in mice (*25*). While AZD2327 is not commercially available it is structurally similar to SNC80, a commercially available δOR-selective agonist (**Fig. 1a**). SNC80 is a known super-recruiter of β-arrestin 2 (*14*) (**Fig. 1b**) and similar to AZD2327 exhibits anxiolytic-like effects in rodents (*17, 20*). These previous findings led us to hypothesize that β-arrestin 2 may be required for the anxiolytic effects of SNC80 and AZD2327. Using two models of anxiety-like behavior, the elevated plus maze (EPM) test and dark-light box transition test (**Fig. 1c**), we measured the behavioral effects of SNC80 in β-arrestin 2 KO mice at a dose known to produce anxiolytic-like effects in wild-type (WT) mice (*20*). As expected, systemic administration of SNC80 (20 mg/kg, s.c.) significantly increased the time WT mice spent in the open arm of the elevated plus maze (**Fig. 1d**; see **Table S1** for two-way ANOVA and post-hoc multiple comparison) and the light chamber of dark-light transition box (**Fig. 1e**; see **Table S1** for two-way ANOVA and post-hoc multiple comparison). As we predicted, the anxiolytic effects of SNC80 were attenuated in β-arrestin 2 KO mice (**Fig. 1d and e**; see **Table S1** for two-way ANOVA and post-hoc multiple comparison). Although the total movement in the elevated plus maze was slightly lower in β-arrestin 2 KO mice than WT mice, no drug effects were observed in both genotypes (**Fig. 1f**; see **Table S1** for two-way ANOVA and post-hoc multiple comparison). Likewise, no statistical difference in total transition was observed in the dark light transition box test (**Fig. 1g** see **Table S1** for two-way ANOVA and post-hoc multiple comparison); however, as previously described, SNC80 produced hyperlocomotive behavior in mice ((*14*), **Fig. S1**, see **Table S1** for detailed statistics).

**Figure 1.**
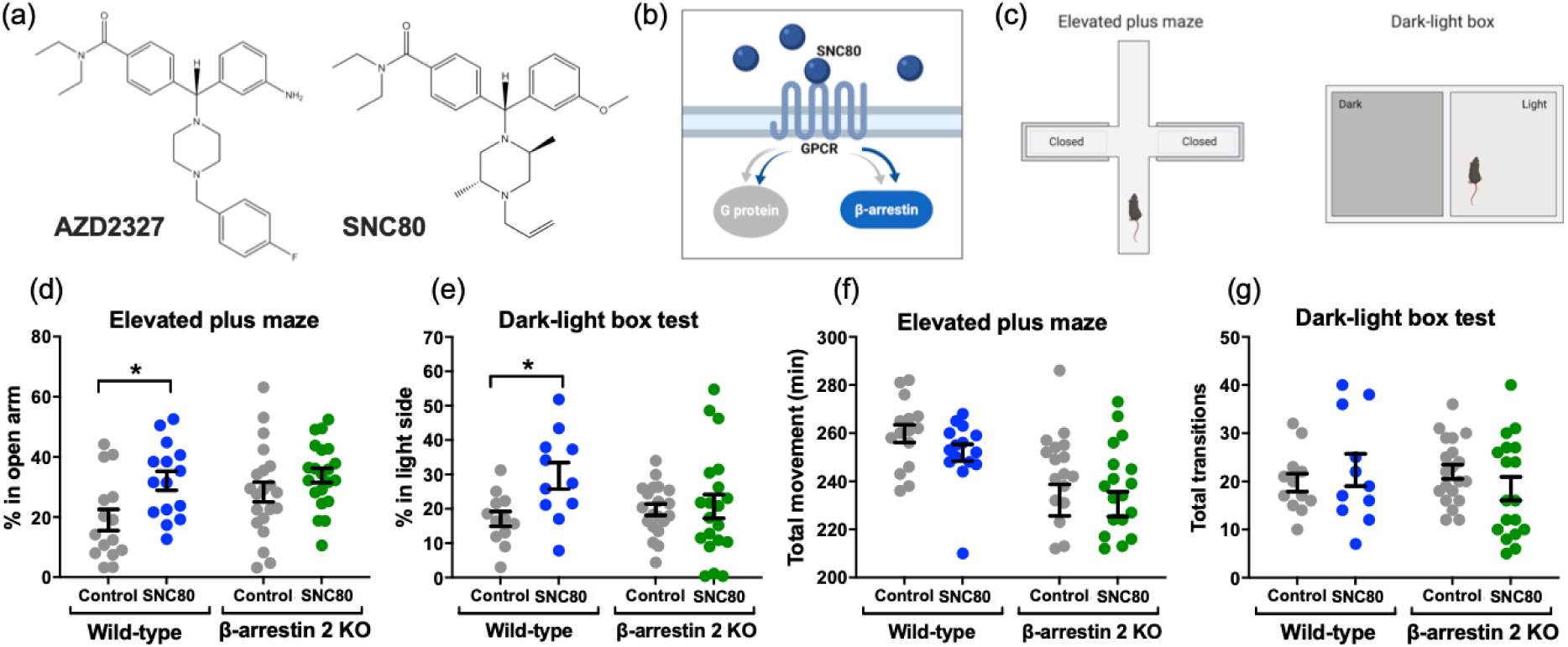
Beneficial role for β-arrestin 2 in reducing anxiety-like behavior. **(a)** Structural similarity between AZD2327, a δOR agonist used in phase II clinical trials for anxious major depressive disorder and SNC80. **(b)** Scheme highlighting that SNC80 is δOR selective agonist, that activates G_i_ proteins but also strongly recruits β-arrestin 2. **(c)** Schematic diagram of the elevated plus maze test and dark-light box test. **(d)** Systemic administration of β-arrestin-biased δOR agonist, SNC80 (20 mg/kg, s.c.), 30 minutes prior to the testing significantly increased percentage of time spent in open arms in WT mice (Control: n=15, SNC80: n=15) but not β-arrestin 2 KO mice (Control: n=21, SNC80: n=21). **(e)** SNC80 increased percentage of time spent in light box in WT mice (Control: n=12, SNC80: n=11), but not in β-arrestin 2 KO mice (Control: n=20, SNC80: n=20). **(f)** The total time of movements were equal between drug treatments. **(g)** No statistical significance was observed in the total transitions between light and dark chambers. (Significance was calculated by two-way ANOVA followed by a Sidak’s multiple comparison; *p<0.05; all values are shown as individaul data points ± S.E.M.).

### The β-arrestin recruiting δOR agonist SNC80 strongly activates ERK1/2 in vitro and in vivo

Activation of κOR has been associated with β-arrestin 2-mediated p38 phosphorylation (*9*). To determine if δOR agonism similarly stimulates mitogen-activated protein kinases (MAPKs), we measured p38, JNK, and ERK1/2 activation in Chinese Hamster Ovarian cells stably expressing δOR and β-arrestin 2 (CHO-δOR-βArr2) following stimulation with 10 µM SNC80, a concentration that will fully activate G-protein signaling and induce β-arrestin 2 recruitment (*26*). We found that SNC80 led to a rapid increase in ERK1/2 phosphorylation within 3 minutes in CHO-δOR-βArr2 cells, which lasted until 60 minutes, in agreement with previous δOR-mediated ERK activation in CHO cells (*27*). We did not observe strong activation of p38 and JNK by SNC80 (**Fig. 2a**). The δOR mediated ERK1/2 signaling in these cells were not an artifact of the recombinant overexpression of δOR and β-arrestin 2 in the CHO cells as we observed a similar profile for ERK1/2 activation in NG108-15 neuroblastoma cells endogenously expressing δOR and β-arrestin (*28-30*) (**Fig. 2b**). We similarly found ERK1/2 activation in several mouse brain regions, known to express δORs, including the dorsal hippocampus, the amygdala and the striatum (*31, 32*) (**Fig. 2c**). The SNC80-induced ERK1/2 activation in these regions was confirmed and quantified by the Western blot analysis of flash-frozen tissue punches upon collection (**Fig. 2d**). Here, we observed that SNC80 (20 mg/kg, i.p.) significantly increased ERK1/2 phosphorylation at the 10-minute time-point in all tested brain regions except for the ventral hippocampus of WT mice (**Fig. 2e-i**; see **Table S2** for one-way ANOVA and post-hoc multiple comparison), and these activations returned to basal levels 30 minutes after the SNC80 administration. The SNC80-induced ERK1/2 activation was not observed in the dorsal hippocampus and the amygdala of δOR KO mice (**Fig. S2**), further confirming this unique profile of ERK1/2 activation by SNC80 is mediated by δOR.

**Figure 2.**
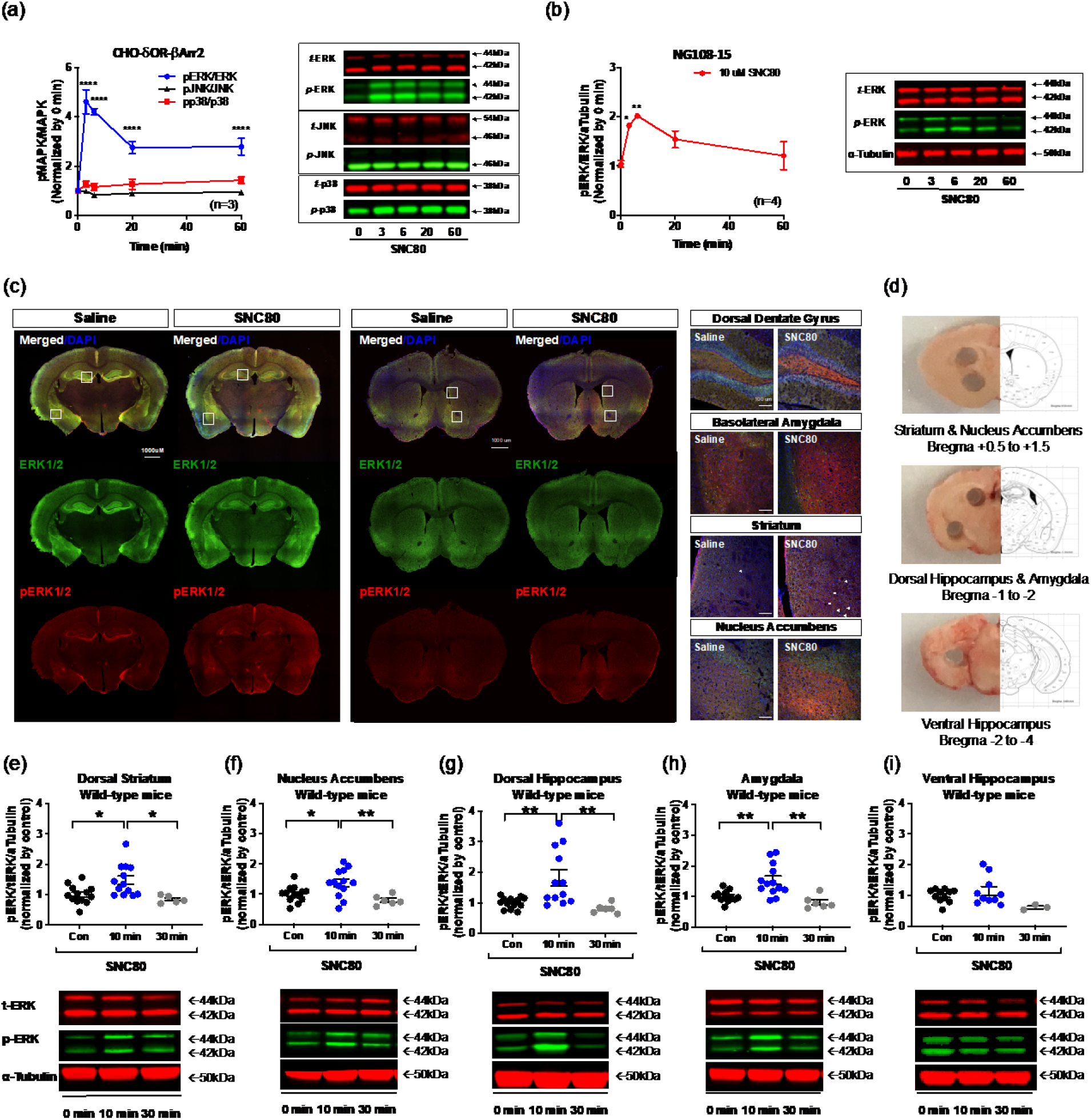
The anxiolytic agonist SNC80 activates ERK1/2 in vitro and in the limbic and striatal regions of the brain. **(a)** 10 uM SNC80 strongly increased ERK1/2 phosphorylation, compared to p38 and JNK, in CHO cells stably expressing δOR and β-arrestin 2. A representative Western blot image is presented to the right. **(b)** SNC80 increased ERK1/2 phosphorylation in a time-dependent manner, in NG-108-15 cells endogenously expressing δOR. A representative Western blot image is shown on the right. **(c)** Representative immunoflouresence staining images of the limbic regions and striatal regions in WT mice. SNC80 (20 mg/kg, i.p.) was administered 10 minutes prior to the perfusion (Green: ERK; Red: pERK; Blue: DAPI). For stitching images (Left), 4x magnification was used to stitch the whole brain slices. Enlarged images (Right) were captured with 20x magnification. Corresponding locations of 20x images were marked in white square of 4x images (Scale bar for 4x represents 1000 μM; Scale bar for 20x represents 100 μM). **(d)** Representative images of brain sections with micropunctures of five specified brain regions for the Western blot analysis. **(e-i)** Activation of ERK1/2 protein activity following systemic adminstration of SNC80 in five wild-type mice brain regions. Representative Western blot images are depicted below the respective bar graph. The number of samples for (g-k) is listed in **Table S2**. (For (a), significance was measured by two-way ANOVA followed by a Dunnett’s multiple comparison. For (b, g-k), significance was analyzed by one-way ANOVA followed by a Tukey’s multiple comparison; *p < 0.05, **p<0.01, ****p < 0.0001; all values are shown as individual data points ± S.E.M.).

### β-arrestin 2 is required to activate ERK1/2 signaling in the limbic structures of the brain

To determine if β-arrestin 2 is responsible for the ERK1/2 activation observed in wild-type mice, we quantified ERK1/2 activation across the same hippocampal, striatal and amygdalar regions of β-arrestin 2 KO mice (**Fig. 3a**) upon systemic administration of SNC80 (20 mg/kg, i.p.) (**Fig. 3b**). While SNC80 still strongly activated ERK1/2 in the nucleus accumbens (**Fig. 3c**; see **Table S2** for one-way ANOVA and post-hoc multiple comparison) and revealed an increasing trend of ERK1/2 activation in striatum (**Fig. 3b**; see **Table S2** for one-way ANOVA and post-hoc multiple comparison) of the β-arrestin 2 KO mice, we did not observe significant SNC80-induced ERK1/2 activation in the dorsal hippocampus, the amygdala, and the dorsal hippocampus of β-arrestin 2 KO mice (**Fig. 3e-g**; see **Table S2** for one-way ANOVA and post-hoc multiple comparison).

**Figure 3.**
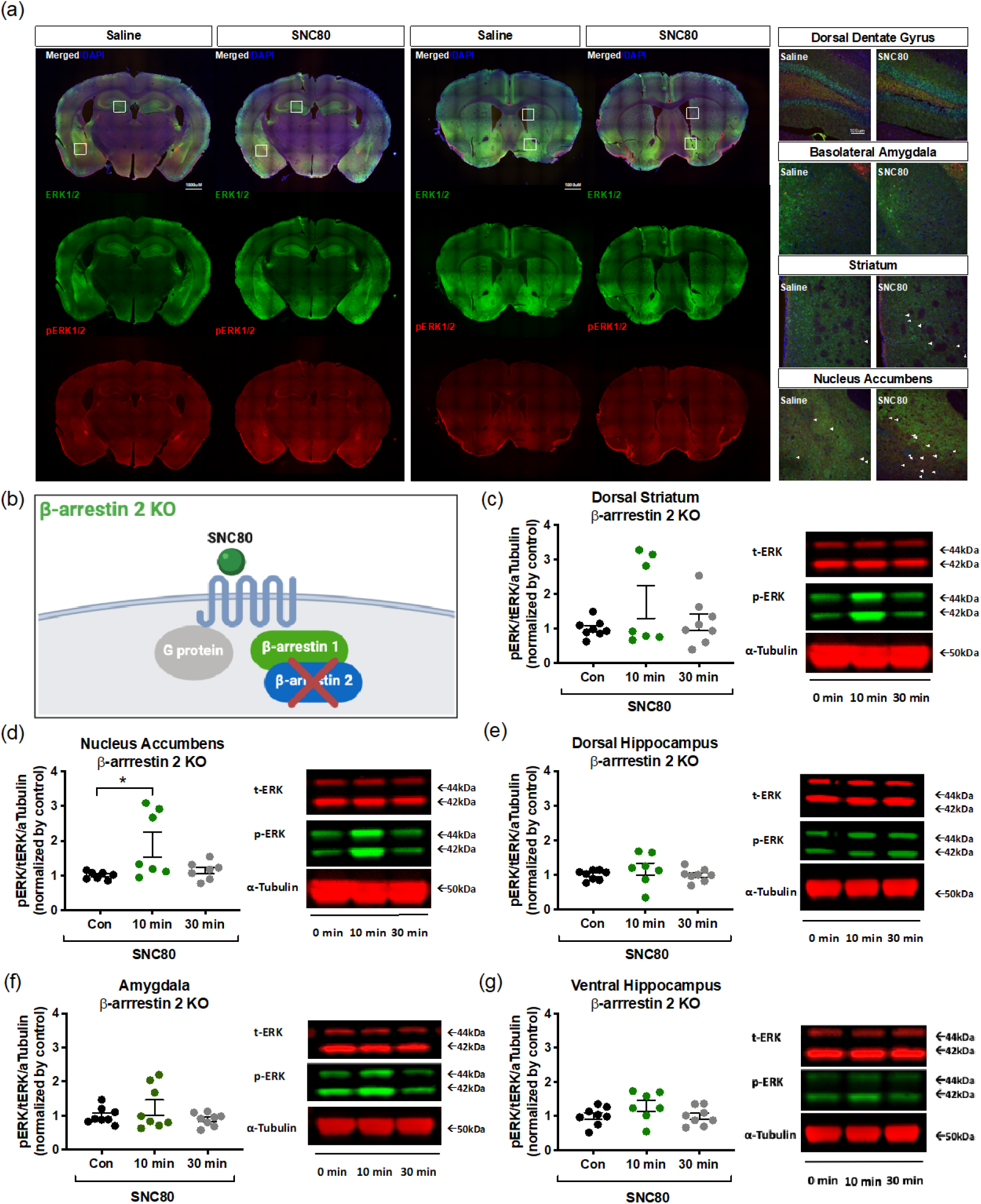
ERK1/2 activation in the amygdala and the dorsal hippocampus are β-arrestin 2 dependent. **(a)** Representative immunoflouresence staining images of the limbic regions and striatal regions in β-arrestin 2 KO mice. SNC80 (20 mg/kg, i.p.) was administered 10 minutes prior to the perfusion (Green: ERK; Red: pERK; Blue: DAPI). For stitching images (Left), 10x magnification was used to stitch the whole brain slices. Enlarged images (Right) were captured with 20x magnification. Corresponding locations of 20x images were marked in white square of 10x images (Scale bar for 10x represents 1000 μM; Scale bar for 20x represents 100 μM). (**b**) Schematic diagram of the cellular context in β-arrestin 2 genetic KO mice. SNC80 (20 mg/kg, i.p.) induced ERK1/2 activation in the dorsal striatum (**c**) and nucleus accumbens (**d**). β-arrestin 2 KO ablated the SNC80-induced ERK1/2 activation in the dorsal hippocampus (**e**) and the amygdala (**f**) with no effects in the ventral hippocampus. **(g)** Representative Western blot images are shown tothe right of the related bar graph. The number of samples for (c-g) is listed in **Table S2**. (Significance was analyzed by one-way ANOVA followed by a Tukey’s multiple comparison; *p < 0.05; all values are shown as individual data points ± S.E.M.).

### ERK1/2 signaling plays a key role in the SNC80-mediated anxiolytic-like effects

We next assessed if the anxiolytic effects of SNC80 were dependent on ERK1/2 activation. We administered wild-type mice with SL327 (50 mg/kg, s.c.), a MEK1/2 inhibitor that indirectly prevents ERK1/2 activation, (**Fig. 4a**) (*33*). We found that pre-administration of SL327 ablated the anxiolytic-like effects of SNC80 (20 mg/kg, i.p.) in WT mice (**Fig. 4b**; see **Table S3** for one-way ANOVA and post-hoc multiple comparison). The hippocampus is a brain region associated with anxiety-like behavior in the elevated plus maze (*34*) and δOR-agonism in the dorsal hippocampus, and the amygdala is associated with reduced anxiety-like behavior in the open field test (*18, 35*). These published findings agree with our observation of SNC80-induced β-arrestin 2-dependent ERK1/2 activity specifically in these two brain regions (**Fig. 3e,f**). Therefore, if the β-arrestin-mediated ERK1/2 signaling in these two regions was critical for the anxiolytic-like effects of SNC80, we would expect ERK1/2 activity to be abolished in these regions in the mice with SNC80 (20 mg/kg, i.p.) and SL327 (50 mg/kg, s.c.). Indeed, we found that SL327 effectively decreased SNC80-induced ERK1/2 activity in the dorsal hippocampus and the amygdala (**Fig. 4c,d**, see **Table S3** for detailed statistics), but not in the dorsal striatum, the nucleus accumbens, and the ventral hippocampus (**Fig. S3**, see **Table S3** for detailed statistics). As an additional approach to investigate the importance of β-arrestin 2 signaling on ERK1/2 signaling and anxiety-like behavior, we utilized a δOR agonist that is a weaker β-arrestin recruiter than SNC80. Specifically, we utilized ADL5859, which affinity (*K*_i_ = 0.84 nM) and G-protein potency (EC_50_=20 nM) at δOR is not significantly different from SNC80 (*K*_i_ = 1.2 nM, EC_50_=10 nM, (*36*)), but has 50-fold lower potency (pEC_50_=6.6±0.1, **Fig. S4a**) and slightly lower efficacy (E_max_ = 119% ±5, relative to Leu-enkephalin **Fig. S4a**) than SNC80 (pEC_50_=8.2±0.1, E_max_ =142%±9, (*14*)). Systemic administration of ADL5859 (30 mg/kg, p.o.) at a dose that produces robust δOR-mediated analgesia (*37*) did not result in ERK1/2 activation in the dorsal hippocampus and the amygdala (**Fig. S4b,c**) nor reduce anxiety-like behavior (**Fig. S4d**), suggesting that ERK1/2 activation in the dorsal hippocampus and the amygdala as well as δOR-mediated anxiolytic effects require a strong recruitment of β-arrestin 2.

**Figure 4.**
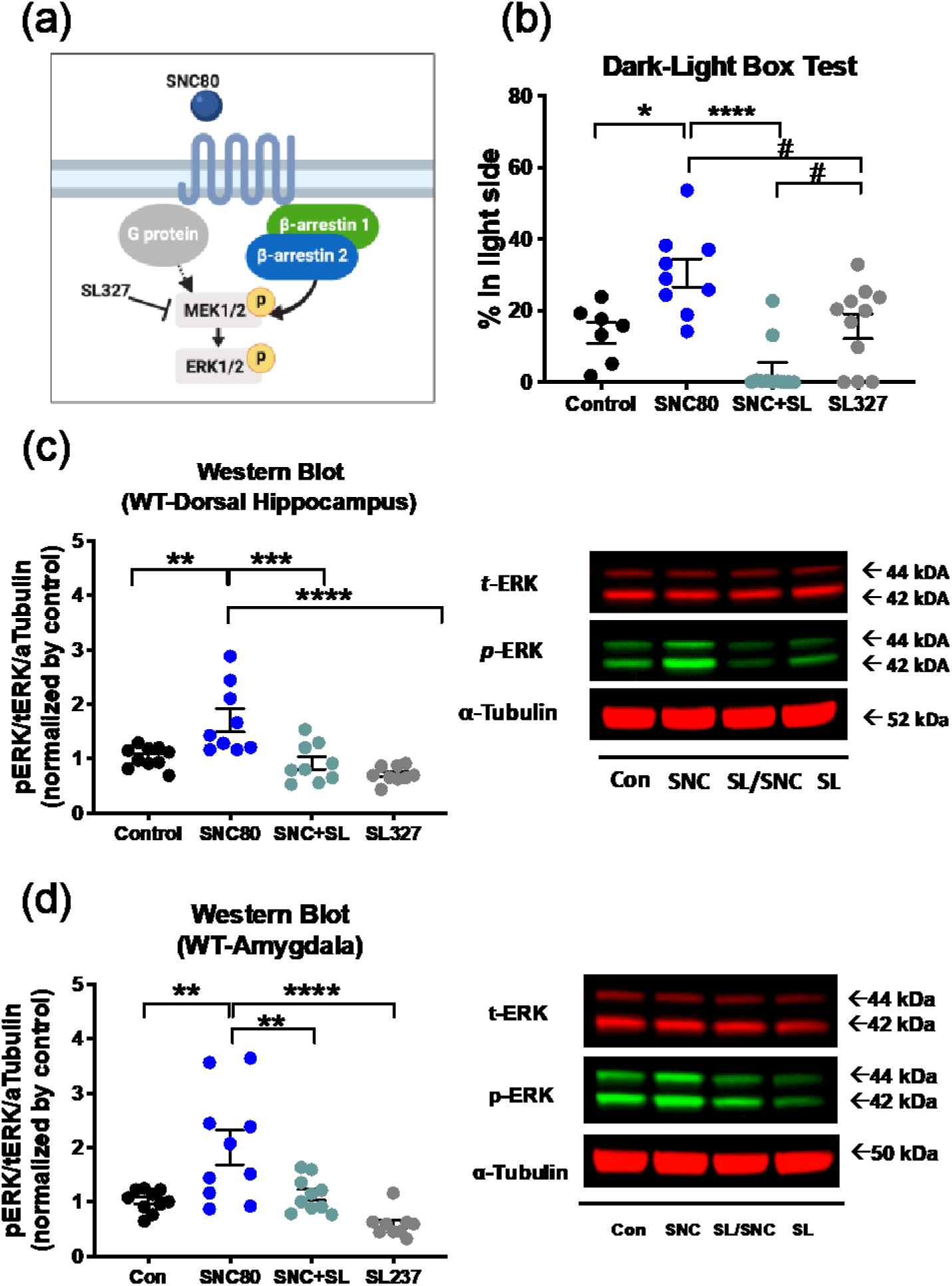
ERK1/2 is required for the anxiolytic-like effects induced by SNC80. **(a)** A schematic diagram of SL327 induced inhibition of SNC80 mediated ERK1/2 signaling. **(b)** SL327 (50 mg/kg, s.c.), which was administered 60 minutes prior to the testing, attenuated the anxiolytic-like effects of SNC80 (20 mg/kg i.p.) administered 30 minutes prior to the testing in WT mice (control: n=8, SNC80: n=12, SNC+SL: n=12, SL327: n=12). **(c-d)** SL327 significantly inhibited SNC80-induced ERK1/2 phosphorylation in the dorsal hippocam and the amygdala. SL327 was also administered 60 minutes prior to the testing, and SNC80 was administered 10 minutes prior to the tissue collection. The number of samples for (c-d) is listed in **Table S3**. (Significance was calculated by one-way ANOVA followed by a Sidak’s or Tukey’s multiple comparison; *p<0.05, **p<0.01, ***p<0.001, ****p<0.0001, #p<0.05; all values are shown as individual data points ± S.E.M.; SNC+SL means SNC80+SL327 and SL means SL327).

### Fear-potentiated startle behavior is correlated with ERK1/2 activity but is not mediated by β-arrestin 2

Besides reducing anxiety-like behavior, δOR activation can also alleviate conditioned fear-related behavior (*17, 38*). Based on our results and previous studies, we hypothesized that SNC80 may similarly reduce fear-related behavior through a mechanism that involves β-arrestin 2. To measure conditioned fear, we utilized a mouse behavior paradigm of fear-potentiated startle (FPS) (**Fig. 5a**). Mice groups were counterbalanced, such that no significant differences between startle reflexes were observed between groups (**Fig. S5**). In WT mice (**Fig. 5b**), we noted that SNC80 (20 mg/kg, i.p.) significantly reduced startle responses to the unconditioned ‘noise’ cue as well as to the conditioned ‘light+noise’ cue (**Fig. 5c**; see **Table S4** for two-way ANOVA and post-hoc multiple comparison). The reduction produced by ‘light+noise’ is larger than the reduction produced by ‘noise’, and thus these reductions result in a significant reduction in % FPS response (**Fig. 5d**; p-value is indicated in figure legends). We also measured if SNC80 would alter %FPS at a dose of 10 mg/kg (i.p.), as this dose of SNC80 has been shown to be effective in other studies effect (*17, 95*), but in our hands we did not observe any significant effects (**Fig. S6**). To our surprise, we found that SNC80 was equally effective in reducing % FPS responses in β-arrestin 2 KO mice (**Fig. 5e,f**; see **Table S4** for two-way ANOVA and post-hoc multiple comparison; **Fig. 5g**; p-value is indicated in figure legends). While SNC80 is a very efficacious recruiter of β-arrestin, it still also fully activates G_i_ protein signaling (*14, 26*). Thus, we next hypothesized that the observed fear-reducing effects of SNC80 could be mediated through G_i_ protein signaling. To address this hypothesis, we utilized a δOR selective agonist, TAN67, which is a very poor recruiter of β-arrestin, and considered G_i_ protein-biased (*14, 26*). However, when we administered TAN67 (25 mg/kg, i.p.) to our WT mice (**Fig. 5h**), TAN67 did not significantly change startle to the ‘noise’ or to the ‘light+noise’ stimuli (**Fig. 5i**; see **Table S4** for two-way ANOVA). However, there was a significant increase in %FPS (**Fig. 5j**; p-value is indicated in figure legends). The lack of a direct effect of TAN67 on noise-alone startle is in agreement with our previous finding that TAN67 did not change basal anxiety-like behavior in the elevated plus maze and dark-light transition test (*20*). Our Western blot analysis of ERK1/2 activities in WT mice revealed that TAN67, decreased ERK1/2 phosphorylation in the dorsal striatum, the nucleus accumbens, the dorsal hippocampus, and the amygdala (**Fig. 6a-d**). In the ventral hippocampus, TAN67 did not alter ERK1/2 activity (**Fig. 6e**), which was similar to the lack of ERK1/2 modulation by SNC80 in this region (**Fig. 2i**) and may be indicative of low δOR expression in the ventral region as previously described (*39*).

**Figure 5.**
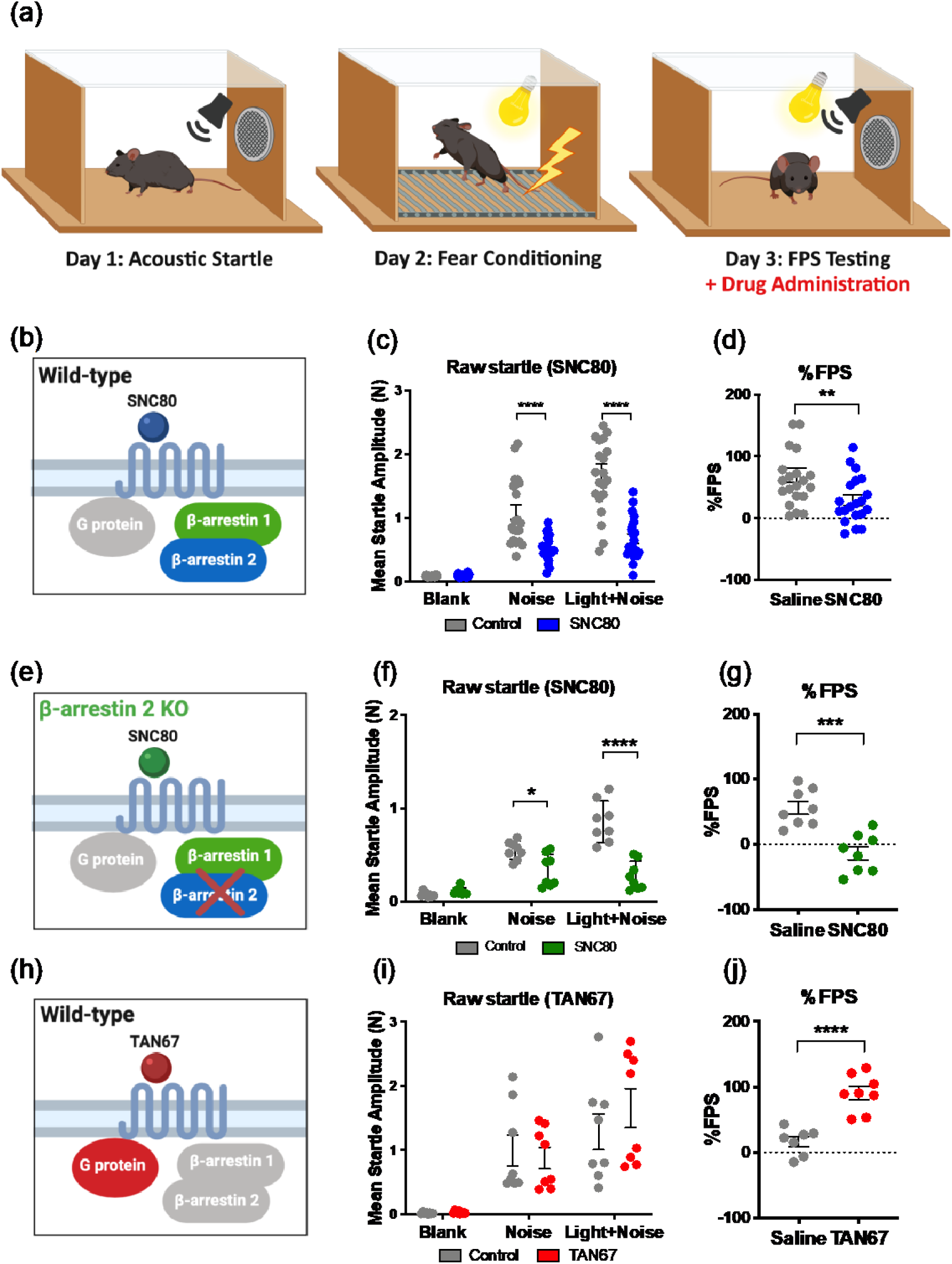
Unique modulation of fear-potentiated startle (FPS) by a β-arrestin-biased and G protein-biased δOR agonist. **(a)** Schematic representation of the three-day experimental paradigm of the fear potentiated startle test; drugs were administered prior to the tests on the third day (See **Figure S5** for Day 1 acoustic startle test). SNC80 (20 mg/kg, i.p.) was adminsitered 30 minutes prior to the testing. **(b-d)** SNC80 (control: n=21, SNC80: n=21) reduced raw startle amplitudes to noise or light+noise and % FPS during the FPS test in WT mice. **(e-g)** SNC80 also reduced raw startle amplitudes to noise or light+noise and % FPS in β-arrestin 2 KO mice (control: n=8, SNC80: n=8). **(h-j)** Yet, G protein-biased agonist, TAN67, increased % FPS in WT mice (control: n=8, SNC80: n=8). (Significance was measured by two-way ANOVA followed by a Sidak’s multiple comparison for (c, f, i) or unpaired t-test for (d, g, j). For (d) *p*=0.0085; for (g) *p*=0.0003; for (j) *p*<0.0001; *p < 0.05, **p<0.01, ***p < 0.001, ****p < 0.0001. All values are shown as individual data points ± S.E.M.).

**Figure 6.**
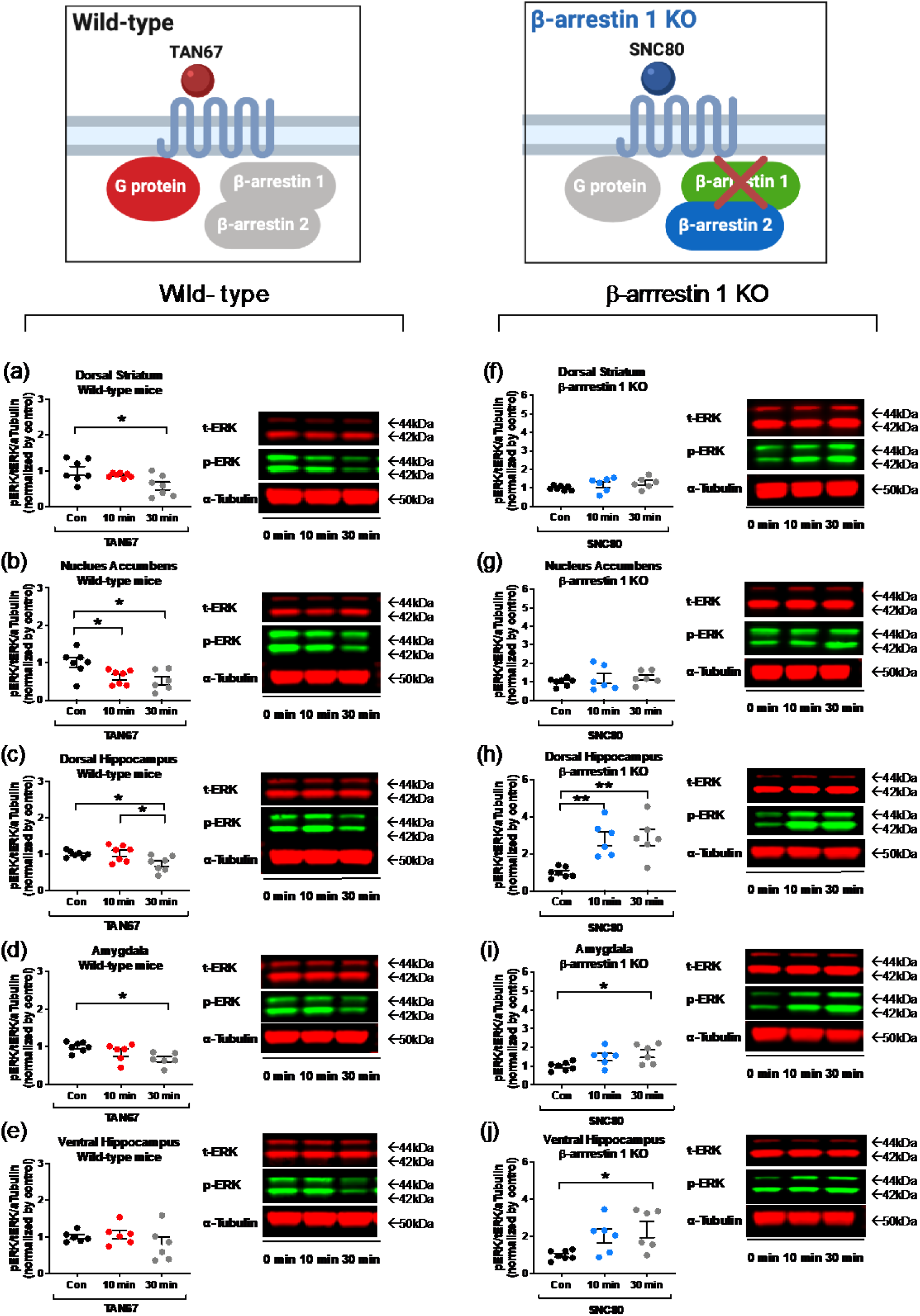
Differential roles for G protein and β-arrestin 1 in δOR agonist-induced ERK1/2 activation. **(a-e)** The G protein-biased δOR agonist TAN67 (25 mg/kg, i.p.), decreased the basal ERK1/2 activity in all tested brain regions of wild-type mice. Representative Western blot images are depicted next to each related bar graph. **(f-j)**. Systemic administration of SNC80 (20 mg/kg, i.p.) did not activate ERK1/2 in the striatal regions of β-arrestin 1 KO mice but resulted in persistent ERK1/2 activation in the dorsal and ventral hippocampus and amygdala. The number of samples is listed in **Table S5**. (Significance was analyzed by one-way ANOVA followed by a Tukey’s multiple comparison; *p < 0.05, **p<0.01; all values are shown as individual data points ± S.E.M.).

### Potential roles for ERK1/2 and β-arrestin 1 in the modulation of conditioned-fear behavior

Our results suggest that SNC80 reduces conditioned fear through a mechanism that does not involve β-arrestin 2 or G protein signaling. Therefore, we next hypothesized that the effect may be mediated by β-arrestin 1 instead. SNC80 is a potent and efficacious recruiter of β-arrestin 1 in vitro (pEC_50_ = 7.9 ± 0.2, E_max_ = 86% ± 5), particularly compared to TAN-67, which recruits β-arrestin 1 with much lower efficacy (pEC_50_ = 7.7 ± 0.4, E_max_ = 14% ± 2) (**Fig. S7**). However, SNC80 is known to induce severe seizures in β-arrestin 1 KO mice (*16*), preventing us from testing the FPS response of SNC80 in this strain. Instead, we measured SNC80-induced ERK1/2 phosphorylation in the β-arrestin 1 KO mice. SNC80 (20 mg/kg, i.p.) was systemically administered at indicated timepoints. In comparison to WT mice (**Fig. 2e-i**), genetic knockout of β-arrestin 1 prevented SNC80-mediated ERK1/2 phosphorylation in the dorsal striatum and nucleus accumbens (**Fig. 6f-g**; see **Table S5** for one-way ANOVA and post-hoc multiple comparison). In the amygdala and dorsal hippocampus of β-arrestin 1 KO mice, SNC80 did increase ERK1/2 phosphorylation (**Fig. 6h-i**), but in contrast to the response observed in WT mice, the activation was sustained for at least 30 minutes. Additionally, we observed ERK1/2 activation in the ventral hippocampus (**Fig. 6j**), a region where we had observed no δOR agonist-mediated modulation of ERK1/2 in WT mice (**Fig. 2i** and **Fig. 6e**), suggesting this pattern of ERK1/2 activity may not be δOR-mediated and is most likely a result of seizure activity.

## Discussion

Here, we investigated the hypothesis that β-arrestin can modulate anxiety- and conditioned fear-related behavior through downstream MAPK activation. By utilizing G protein- and β-arrestin-biased δOR agonists together with β-arrestin-isoform selective knockout mice, we discovered that G protein, β-arrestin 1, and β-arrestin 2 uniquely modulated ERK1/2 activity resulting in differential outcomes in mouse models of anxiety/fear-related behavior. Our results suggest that the reduction in anxiety-like behavior by SNC80 required the presence of β-arrestin 2 as well as activation of ERK1/2. Distinctly in the dorsal hippocampus, we found that ERK1/2 activation was β-arrestin 2-dependent (**Fig. 7a**). We also found that SNC80-induced ERK1/2 activation in the nucleus accumbens and the dorsal striatum required β-arrestin 1, which may be part of the mechanism by which SNC80 decreased FPS (**Fig. 7b**) as well as increase general locomotion (**Fig. S1**). Notably, G protein-biased signaling by TAN67 reduced ERK1/2 phosphorylation and was correlated with increased FPS (**Fig. 7c**).

**Figure 7.**
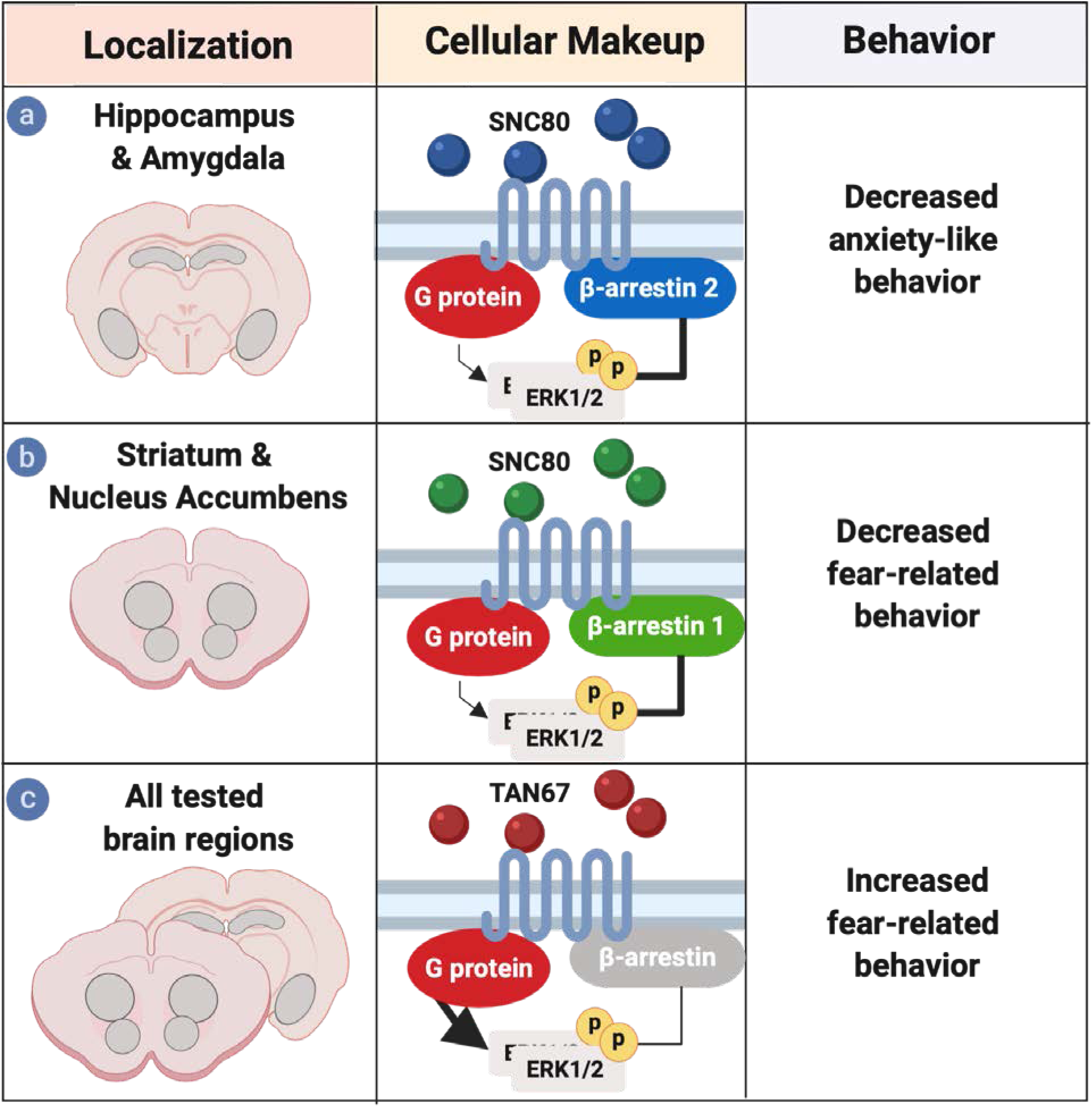
Graphical summary. δOR agonists differentially induce ERK activation in a β-arrestin isoform specific manner to modulate anxiety- and conditioned fear-related behaviors. SNC80 induced β-arrestin 2-mediated ERK1/2 activation (Cellular Makeup) in the hippocampus and amygdala (Localization) decreased anxiety-like behavior **(a)**, whereas an increase in conditioned fear-related behavior (Behavior) by TAN67 can be linked to decreased ERK1/2 activity in all tested brain regions except for the ventral hippocampus **(b)**. Decreased FPS could not be correlated with G protein or β-arrestin 2 signaling, but may involve β-arrestin 1-dependent ERK1/2 signaling in the dorsal striatum and nucleus accumbens **(c)**.

### Differential roles for β-arrestin isoforms in neuropsychiatric behavior

The β-arrestin proteins were discovered in quick succession (*40, 41*), surprisingly, despite the availability of genetic KO mice for each isoform (*42, 43*). Studies investigating β-arrestin in the CNS have largely focused on β-arrestin 2 and have generally neglected β-arrestin 1 (*44, 45*). A potential reason for the preference of studying β-arrestin 2 may be that when the β-arrestin 2 knockout mice were generated their first utilization was to highlight the proteins’ involvement with the CNS-mediated adverse effects of µ-opioid receptor agonism (*46, 47*). Researchers have only recently begun to utilize β-arrestin 1 KO mice to study various neurological disorders. In 2016, Pradhan *et al*. found that different δOR agonists either preferentially recruited β-arrestin 1 leading to δOR degradation or recruited β-arrestin 2 causing δOR resensitization (*15*). A study in 2017 found that amphetamine-induced hyperlocomotion was amplified in β-arrestin 1 KO mice, but attenuated in β-arrestin 2 KO mice (*48*), further emphasizing the importance of studying both β-arrestin isoforms. While our study identified that SNC80-induced anxiolytic-like effects were β-arrestin 2-dependent, β-arrestin 2 KO mice still exhibited the fear-reducing effect of SNC80, which could suggest a potential role for β-arrestin 1 in the modulation of fear-related behavior. Particularly, we noted that β-arrestin 1 KO abolished SNC80-induced ERK1/2 activation in the striatum and nucleus accumbens. Our findings were in line with the extensive *in-situ* hybridization studies on differential β-arrestin isoform levels in neonatal and postnatal rats (*49, 50*) and studies showing relatively high β-arrestin 1 and low β-arrestin 2 expressions in the striatal regions (*41, 49, 51*). In contrast, SNC80-induced ERK1/2 activation in the dorsal hippocampus and amygdala was β-arrestin 2-dependent, which agreed with reports of stronger expression of this isoform in those areas (*41, 51*). Unfortunately, we were limited in our ability to assess whether SNC80-induced reduction in FPS would be attenuated in β-arrestin 1 KO mice, as SNC80 produces severe seizures in these mice (*16*), a phenomenon we have also observed ourselves and found to be accompanied by strong and persistent ERK1/2 activation in the dorsal hippocampus. Further investigation of the roles of β-arrestin 1 in neuropsychiatric behavior may be feasible using a conditional knockout approach; currently conditional β-arrestin 2 knockout mice already exist (*52*), but conditional β-arrestin 1 knockout mice have not yet been reported.

### A unique role for G_i_ protein signaling and ERK1/2 signaling in conditioned fear-related behavior

In contrast to the β-arrestin-mediated activation of ERK1/2, we found that selective G protein signaling at the δOR by TAN67 decreased ERK1/2 activation (**Fig. 6a-d**) and was associated with increased FPS (**Fig. 5j**). TAN67 is a known weak recruiter of β-arrestin 1 and 2 (*14*) (**Fig. S7**). This result is in agreement with the observation and that blocking G_i/o_ protein signaling using pertussis toxin in the basolateral amygdala reduced FPS (*53*) and parallels finding that TAN67 did not reduce unconditioned anxiety-like behavior in naïve mice (*20*). Additionally, a study in ovariectomized mice found that estradiol benzoate, an anxiogenic estradiol prodrug (*54*), decreased ERK in the hippocampus (*55*), which supports our finding that decreased ERK1/2 is correlated with increased fear. It is noteworthy that both G_i_ protein and β-arrestin can activate ERK1/2 albeit through different mechanisms (*56, 57*). One explanation for our observation is that TAN67 competes with the endogenous δOR agonist Leu-enkephalin, which is a much more efficacious recruiter of β-arrestin (*14*) (**Fig. S8**). Based on observations of enhanced anxiety-like behavior in preproenkephalin KO mice, Leu-enkephalin has anxiolytic-like effects by itself (*58, 59*), which is in line with reports that the δOR antagonist, naltrindole, is anxiogenic (*60*). As a weak β-arrestin recruiter, TAN67 would attenuate any baseline Leu-enkephalin-induced β-arrestin-mediated ERK activity (**Fig. S8**). Because ADL5859 has a low potency for β-arrestin 2 recruitment, the same pharmacological principle described above holds true, with the exception that at supersaturating doses, ADL5859 will begin to efficaciously recruit β-arrestin 2, whereas TAN-67 will not. In contrast, because SNC80 is a stronger β-arrestin recruiter than Leu-enkephalin, it will elevate basal ERK1/2 activity produced by endogenous opioids. This hypothesis would also explain why SNC80-induced ERK1/2 activation is quite variable and produces on average only a two-fold increase in pERK1/2 levels.

### β-arrestin serves as a scaffold for a range of kinases and effectors

In our study, we found that δOR agonism strongly activates ERK1/2 compared to p38 and JNK, and that the ERK1/2 activity induced by SNC80 was negatively correlated with FPS. Other GPCRs, besides the δOR, may also require β-arrestin-dependent ERK1/2 signaling for modulation of fear. For example, in the infralimbic prefrontal cortex, β-adrenergic receptor activation can promote the extinction of contextual fear memory (*61*). In a recent study, it was shown that in this same brain region β-arrestin 2-dependent ERK1/2 activation was required for β-adrenergic receptors agonists to stimulate extinction learning of cocaine-induced reward memories (*52*). β-arrestin 2-mediated signaling in the CNS is not exclusive to ERK1/2 signaling; following κOR activation, β-arrestin 2 can scaffold with p38 as part of a potential mechanism for the aversive effects of κOR agonists (*62*). A β-arrestin 2 scaffold of AKT, GSK3β and PP2A has been proposed as a mechanism for stabilizing mood (*63*), highlighting that β-arrestin signaling is also not limited to MAPKs. In fact, because β-arrestin 1, in contrast to β-arrestin 2, contains a nuclear translocation sequence (*64*), it can translocate to the nucleus and regulate gene transcription (*65, 66*). While in this study we report δORs require β-arrestin-dependent ERK signaling for reduction in anxiety-like behavior, it is certainly possible that other GPCRs may engage different intracellular signaling pathways following β-arrestin recruitment.

### Fear- and anxiety-like behaviors rely on shared but distinct neural circuits

In our study, we observed that β-arrestin 2-dependent ERK1/2 activity in the dorsal hippocampus was associated with reduced anxiety-like behavior. Generally, CA1 regions of the ventral hippocampus are associated with responding to contextually-conditioned anxiogenic stimuli than dorsal CA1 regions (*67, 68*), whereas the dorsal hippocampus is involved with cognitive functions, including exploration, navigation and memory (*69*). Still, our finding that δOR signaling in the dorsal hippocampus is connected to the anxiolytic-like effects of SNC80, agrees with a study showing that intra-dorsal CA1 injection of the δOR antagonist naltrindole is anxiogenic (*35*). Additionally, the anxiolytic-like effects of SNC80 may also involve β-arrestin 2-dependent ERK1/2 signaling in the amygdala, a region more commonly associated with innate anxiety-like behavior (*70, 71*). The basolateral amygdala (BLA) also plays an important role in fear conditioning (*72*), including FPS (*73*). The BLA receives dopaminergic inputs from the ventral tegmental area and projects to the nucleus accumbens. It has been proposed that dopaminergic signaling in the BLA is important for cue-dependent fear-conditioning, such as FPS (*74*) and that the BLA to nucleus accumbens projection is critical for consolidation of memories associated with aversive effects such as foot shock (*75, 76*). Future studies using conditional β-arrestin and/or δOR KO mice could help assess to what degree dopamine receptor-mediated β-arrestin signaling is required for the observed behaviors.

Our finding that ERK1/2 activation in the striatal regions was ablated in β-arrestin 1 KO mice may suggest that striatal β-arrestin 1-mediated ERK1/2 signaling is critical for the modulation of the expression of conditioned fear-related behavior. Processing and executing emotional behaviors in tasks such as the elevated plus maze and FPS tests engages multiple overlapping, yet distinct, brain regions and circuits involved in memory retrieval, locomotion, decision making, reward, and mood (*72*). Further studies with circuit-based approaches are necessary to assess the role of biased signaling pathways in the acquisition and expression of conditioned fear-related behavior.

### Beneficial roles of β-arrestin signaling

For the longest time, β-arrestin 2 has been associated solely with adverse effects of opioid activation, including tolerance, constipation, respiratory depression, aversion and alcohol use (*20, 46, 77*). These studies fueled a drive to develop G protein-biased opioids to treat pain and other disorders with an improved therapeutic window (*78, 79*). Yet, recently a number of studies have started to push back against this narrative (*80-84*). Clearly, β-arrestin signaling is not inherently negative as the therapeutic effects of lithium and fluoxetine and D_2_R agonists seem to depend on β-arrestin 2 (*85-87*). The increased propensity for β-arrestin 1 KO mice to experience SNC80-induced seizure points to a potential beneficial role for this isoform in maintaining seizure threshold, which could be of use in the treatment of epilepsy. In this study, we provide additional insights regarding potential therapeutic benefits of β-arrestin signaling in reducing anxiety-like behavior. Providing adequate relief of chronic pain is not trivial, partly because it is often associated with negative affect (*88, 89*) including anxiety, which may exacerbate pain (*90*). δOR agonists have been proposed as potential treatment for chronic pain disorders (*91*), partly because they have the ability to not only provide analgesia, but also treat comorbid anxiety and depression (*17, 20, 92*). However, our results would argue that developing G protein-biased δOR agonists may produce drugs that are suboptimal for the treatment of complex chronic pain; our findings suggest such a drug would not alleviate co-morbid fear and anxiety, but potentially even worsen these symptoms. Thus, our study results argue in favor of a reassessment of drug development efforts that seek solely to identify G protein-biased drugs. Instead, we propose that efforts should be directed towards the development of drugs with finely tuned bias and, if possible, towards development of molecules that are biased against a single β-arrestin isoform rather than both isoforms and are circuit-specific.

### Conclusion

Overall, our results begin to reveal the complex- and context-specific nature of GPCR biased signaling in modulation of fear-related and anxiety-like behavior. These results expand our current understanding of therapeutic effects of β-arrestin signaling in mood disorders, which ultimately may aid development of more efficacious pharmacological treatment options for these disorders.

## Materials and Methods

### Animals

Wild-type (WT) C57BL/6 male mice were purchased from Envigo (Indianapolis, IN), and β-arrestin 1 or 2 global knockout (KO) mice as well as δOR KO mice were bred in our facility (*14, 93, 94*). The knockout mice have been rederived every three years by crossing the KO mice with WT C57BL/6 mice from Envigo to reduce genetic drift. Adult mice (8-10 weeks, 25 ± 3g) were group housed (3-5 mice) in a single ventilated Plexiglas cage. Mice were maintained at ambient temperature (21°C) in an animal housing facility approved by the Association for Assessment and Accreditation of Laboratory Animal Care and animals were kept on a reversed 12-hour dark-light cycle (lights off at 10:00, lights on at 22:00). Food and water were provided ad libitum. Purchased mice were acclimated for one week prior to the experiments. All animal protocols (#1305000864 by RMvR) were preapproved by Purdue Animal Care and Use Committee and were in accordance with the National Institutes of Health’s Guide for the Care and Use of Laboratory Animals.

### Drug preparation and administration

SNC80 (#076410, Tocris, Thermo Fisher Scientific, Waltham, MA) was diluted in slightly acidic saline pH5-6. TAN67 (#092110, Tocris, Thermo Fisher Scientific) was diluted in sterile saline. SL327 (#19691, Tocris, Thermo Fisher Scientific) was diluted in 5% DMSO, 10% Cremophore (Millipore Sigma, Burlington, MA) and 85% saline. ADL5859 (#1751, Axon Medchem, Reston, VA) was diluted in 0.5 % methylcellulose and 0.1 % tween 80 as indicated in (*37*). To investigate our hypotheses, we utilized several δOR agonists: SNC80, TAN67, and ADL5859. Doses for each drug were based on those that proved to impact behavior through δOR (*14, 20, 37*). For SNC80, we used a dose of 20 mg/kg (*14, 20*). We have also tested 10 mg/kg SNC80 (i.p.). While this dose was effective in other studies effect (*17, 95*), in our hands, we did not observe any significant effects (**Fig. S6**). TAN67 was tested at a dose of 25 mg/kg (*14, 20*), whereas ADL5859 was administered at a dose of 30 mg/kg (*37*). We also used the MEK inhibitor SL327 at a 50mg/kg dose to prevent ERK phosphorylation, as this inhibitor has been reported to cross the blood-brain barrier (*96, 97*). Except for ADL5859, which was administered with per oral (p.o.), all drugs were administered either intraperitoneal (i.p.) or subcutaneous (s.c.), and specific times prior to behavior or brain tissue extraction can be found in figure legends. Additionally, separate batches of mice with no prior history of drug injection were used for the brain collection and behavioral tests to test earlier time-points of ERK1/2 signaling (such as 10 minutes).

### Elevated-plus maze test

The elevated-plus maze test was performed as previously described (*20*). Mice were allowed to explore the maze for 5 minutes, and arm entries and time spent in each arm were recorded with a camera positioned above the maze.

### Dark–light transition box test

The test was performed based on previously established protocols (*20, 93*) Testing was conducted without a habituation session to the boxes and a 1/2 area dark insert was placed in the locomotor boxes, leaving the remaining 1/2 of the area lit as described previously (*98*). Two LED lights were inserted above the light portion of the testing chamber where the lux of the light region ranged from 390-540 lumens and dark chamber lux ranged from 0-12 lumens. For testing, animals were placed in the light portion of the chamber and testing began upon animal entry. Time spent in the dark and light chambers as well as their locomotor activity was recorded for 5 minutes with a photobeam-based tracking system.

### Fear potentiated startle (FPS) test

Startle reflexes of mice were recorded in the startle reflex chambers (25.8 × 25 x 26.5 cm) using the Hamilton Kinder Startle Monitor system (Kinder Scientific, Poway, CA). On the conditioning day, all subjects were conditioned with 40 conditioning trials by a fixed 2 minute inter-trial interval (ITI), and FPS responses were tested on the following day. The fear conditioning and FPS parameters were based on a previously established protocol (*99*).

### Preparation of tissue homogenates

After drug injections, mice were euthanized by carbon dioxide asphyxiation and rapidly decapitated. Based on our previous studies, we have particularly chosen carbon dioxide asphyxiation over other euthanasia methods that may potentially increase basal ERK1/2 activity in the brain (*100*). The collected brains were first sliced as coronal sections (1.5-2.0 mm) with a brain matrix (#RBMS-205C, Kent scientific, Torrington, CT), and then flash-frozen in dry-ice-chilled 2-methylbutane (−40 °C) (#03551-4, Fisher Scientific). Regions of interest were collected from these slices using a 1 mm biopsy micropunch (#15110-10, Miltex, Plainsboro, NJ) as follows: dorsal striatum and nucleus accumbens (A/P: +0.5 mm to +1.5 mm), dorsal hippocampus and amygdala (A/P: -1 mm to -2 mm), and ventral hippocampus (A/P: -2 mm to -4 mm) (*101*). The punches targeted a specific region and produced enough tissue to run several blots. However, it is noteworthy that the extracted tissue may contain small amounts of tissue from neighboring regions. For example, while the punches for the amygdala primarily consisted of the BLA, the tissue will also have included a small portion of the central amygdala. Collected tissues were further homogenized with a tissue grinder (#357535 & 357537, DWK Life Sciences, Millville, NJ) in RIPA buffer mixed Halt™ Protease and Phosphatase Inhibitor Cocktail (#1861280, Thermo Fisher Scientific). Samples were further prepared based on previously established protocols (*100*). Data depicted in **Fig. 2e-i** also includes the data collected for SNC80 at the 0 min or 10 min time points in the experiment depicted in **Fig. 4c,d** to represent the full range of observed SNC80 induced ERK1/2 activation in mice tested in separate cohorts at different occasions.

### Cell culture

Chinese hamster ovarian CHO-δOR-βArr2 cells (DiscoverX, Fremont, CA) U2OS-δOR-βArr1 (DiscoverX), and NG-108-15 cells (HB-12317™, ATCC®, Manassas, VA) were cultured as recommended by the manufacturer and maintained at 37° C/5 % CO_2_. Cells were seeded in a clear 6 well plate (Corning™, Thermo Fisher Scientific) with 250,000 cells/2 mL/well. On the following day, all growth media was aspirated and changed into 1 mL serum-free Opti-MEM (#31985070, Gibco®, Thermo Fisher Scientific). The next day, cells were challenged with 10 µM drugs (SNC80) for a specific duration (0, 3, 6, 20, and 60 minutes). All drugs were diluted in Opti-MEM prior to administration. The media was aspirated following the challenge and 100 µL RIPA buffer was added to collect the samples on ice. Using cell scrapers (#353089, Thermo Fisher Scientific), all samples were dislodged from the 6-well plate, collected and stored at -30° C until usage. For the Western blot, the collected samples were quantified with the Bradford assay and samples were prepared with 4 x Laemmli and boiled at 95° C for 5 minutes. The CHO-δOR-βArr2 cells were also used to measure β-arrestin recruitment using the DiscoverX PathHunter Assay as previously described (*14*).

### SDS-Page and Western blot

Samples (20 µL containing 10 µg protein) were loaded per well of a NuPage 4-12 % Bis-Tris gradient gels (#NP0336BOX, Thermo Fisher Scientific), and the SDS-Page gel was subsequently transferred to nitrocellulose membranes (#1620115, BioRad) by the Western blot. Membranes were incubated following previously established protocols (*100*). For reproducibility, detailed information regarding the antibodies used in the study are listed in **Table S6**. Prepared samples were scanned using the LiCor Odyssey® CLx Scanner (Li-Cor, Lincoln, NE). By utilizing the Li-Cor secondary antibodies, we were able to detect the MAPK, pMAPK, and α-Tubulin on the same blot without the need of stripping/reblotting. In the same membrane, each band was cut based on their size. For instance, ERK1/2 and pERK1/2 bands were collected around 42/44 kDa and α-Tubulin band was collected around 50 kDa in the same membrane. For statistical analysis, we normalized the pMAPK/MAPK ratio to α-Tubulin in case drug treatment changed ERK1/2 levels.

### Preparation of tissue for immunofluorescence

To preserve the intact ERK1/2 activity in vivo for fluorescence microscopy, mice were transcardially perfused before isolation of brain tissue. Thirty minutes prior to transcardiac perfusion, mice were administered with 100/10 mg/kg Ketamine/Xylazine. Ten minutes prior to perfusion, 20 mg/kg SNC80 (i.p.) or a corresponding volume of saline was administered to the mice. Mice were then perfused with 30 mL of cold PBS and 4% paraformaldehyde (#100503-916, VWR, Radnor, PA) and were immediately decapitated to collect the brains. The brains were fixated in 4% paraformaldehyde overnight, dehydrated in 30% sucrose, and then embedded in Frozen Section Compound (#3801480, Leica, Wetzlar, Germany). Frozen brains were sliced at a width of 30 μm using the Leica cryostat and permeabilized in 100% methanol at - 20 °C for 10 minutes. The slices were blocked in 5 % Normal goat serum (#S26-100ml, Millipore Sigma) for an hour then stained with primary antibodies as listed in **Table S6**. For immunofluorescence labeling, the sections were incubated in the secondary antibodies according to the previously established protocol (*102*) and as listed in **Table S6**. After the final washing, the nuclei of the sections were stained and the slices were mounted on a glass slide with Vectashield® (#H-1200, Vector lab, Burlingame, CA). Images were acquired with a Nikon confocal microscope and assembled in Adobe Photoshop CS6 (Adobe).

### Statistical analysis

The maximum amplitude of the startle response was measured from the average of all responses for each trial type (12 noise-alone, 12 light+noise) of each mouse. Fear-potentiated startle response was analyzed using raw (maximum) startle amplitudes and proportional changes of each trial type (noise-alone, light+noise), which is shown as % FPS in the graphs. The proportional change score (% FPS) was calculated as follows: (startle response to light and noise – startle response to noise)/startle response to noise x 100. Thus, %FPS is a sensitive measure that adjusts for individual and group differences (for example, possible non-specific effects of drug treatment) in startle response magnitude that may be observed on noise-alone and light + noise trials (*103*).

All data are presented as individual data points (or means) ± standard error of the mean (S.E.M.). Assays with one independent variable were analyzed for statistical significance using a one-way Analysis of Variance (ANOVA), whereas assays with two independent variables were analyzed using a two-way ANOVA. If a significant deviation of the mean was identified, an appropriate post-hoc analysis was performed as indicated in the supplemental table or the corresponding figure legends. Gaussian distribution of our datasets was assessed using the D’augostino and Pearson analysis. We excluded one outlier in our wild-type cohort that received SNC80 and one that received TAN67 in the FPS assay based on the Grubbs’ test (α = 0.05). In the dark-light and elevated plus maze tests we excluded subjects that were frozen/stationary for >95% of the experimental time. All statistical analysis was conducted using GraphPad Prism 7 (GraphPad Software, La Jolla, CA).

## Acknowledgements

We appreciate Dr. Marcus M. Weera for technical assistance with the fear-conditioning experiments. Included diagrams were created with BioRender.

## Author Contributions

MJ.K.: formal analysis, investigation, writing – original draft, writing – review & editing, visualization; T.C.: formal analysis, investigation, writing – review & editing; G.E.M.: formal analysis, investigation, writing – review & editing; A.A.M.: formal analysis, investigation; A.M.G.: formal analysis, investigation, writing – review & editing; A.L.: formal analysis, investigation; J.A.C.: formal analysis, writing – review & editing, supervision; R.M.vR.: Conceptualization, formal analysis, investigation, writing – original draft, writing – review & editing, visualization, supervision, project administration, funding acquisition.

## Data and Materials Availability

All data needed to evaluate the conclusions in the paper are present in the paper or the Supplementary Materials.

