## Supplemental Materials for "β-arrestin-dependent ERK signaling positively correlates with reduced anxiety-like and conditioned fear-related behavior in mice"

**Lists of Supplemental Materials**

**Table S1.** Statistical analysis of anxiety-like behavior upon systemic administration of SNC80 in WT and β-arrestin 2 KO mice

**Table S2.** Statistical analysis of anxiety-like behahvior and ERK1/2 expression levels upon time-series administration of SNC80 in WT and β-arrestin 2 KO mouse brain

**Table S3.** Statistical analysis of ERK1/2 expression levels upon administration of SNC80 in the presence/absence of SL327 in WT mouse brain

**Table S4.** Statistical analysis of fear-related behavior upon systemic administration of SNC80 or TAN67 in WT and β-arrestin 2 KO mice

**Table S5.** Statistical analysis of ERK1/2 expression levels upon time-series administration of TAN67 in WT mousee and SNC80 in β-arrestin 1 KO mouse brain

**Table S6.** Antibody information for the Western blot

**Fig. S1.** Locomotor effects of drug/vehicle treatment in the WT or β-arrestin 2 KO mice in the dark light upon administration of drugs

**Fig. S2.** SNC80 does not activate ERK1/2 in the dorsal hippocampus and the amygdala of δOR KO mice

**Fig. S3.** SNC80-induced ERK1/2 activation is partly affected by SL327 in the striatal regions of the brain

**Fig. S4.** A δOR agonist, ADL5859, does not affect ERK1/2 activity and anxiety-like behaviors of WT mice

**Fig. S5.** Mice groups for FPS tests were counterbalanced based on baseline acoustic startle response

**Fig. S6.** A low dose SNC80 does not affect fear-related behavior of WT mice

**Fig. S7.** β-arrestin 1 recruitment levels by G-protein-biased (TAN67), β-arrestin-biased (SNC80), and non-biased (Leu-Enk) δOR agonist in U2OS-δOR-βArr1 cells

**Fig. S8.** A diagram respresenting the pharmacological competition between two biased agonists and an endnogenous opioid in relations to their ability to modulate ERK1/2 signaling

### Supplemental Materials

**Table S1. Statistical analysis of anxiety-like behavior upon systemic administration of SNC80 in WT and β-arrestin 2 KO mice**

Statistical differences of anxiety-like behaviors in WT or β-arrestin 2 KO mice shown in **Fig. 1**. Significance between groups was calculated by two-way ANOVA followed by a Sidak's multiple comparison (*$p$<0.05, and ns=not significant).

| Subfigure | Behavior test | Genotype | Drug | # of samples | Test | Source of Variation | F-value | p-value | Post hoc analysis | Group Comparison | Mean Diff. | p-value | Significance | - |
|---|---|---|---|---|---|---|---|---|---|---|---|---|---|---|
| Figure 1-d | Elevated plus maze test | WT & βarr2 KO | SNC80 (20 mg/kg, s.c.) | WT-Control: 15<br>WT-SNC80: 15<br>B2-Control: 21<br>B2-SNC80: 21 | Two-Way ANOVA test | Interaction<br>Genotype factor<br>Drug factor | F(1,68) = 1.429<br>F(1,68) = 3.15<br>F(1.68) = 8.781 | 0.236<br>0.0804<br>0.0042 | Sidak's Multiple Comparison Test | WT: Con vs. SNC80 | -13.0400 | 0.0164 | * | |
| | | | | | | | | | | B2 KO: Con vs. SNC80 | -5.5450 | 0.3200 | ns | |
| Figure 1-e | Dark-light box test | WT & βarr2 KO | SNC80 (20 mg/kg, s.c.) | WT-Control: 12<br>WT-SNC80: 11<br>B2-Control: 20<br>B2-SNC80: 20 | Two-Way ANOVA test | Interaction<br>Genotype factor<br>Drug factor | F(1,59) = 3.677<br>F(1,59) = 1.039<br>F(1.59) = 4.978 | 0.06<br>0.3122<br>0.0295 | Sidak's Multiple Comparison Test | WT: Con vs. SNC80 | -12.5200 | 0.0232 | * | |
| | | | | | | | | | | B2 KO: Con vs. SNC80 | -0.9467 | 0.9584 | ns | |
| Figure 1-f | Elevated plus maze test (Total movement - min) | WT & βarr2 KO | SNC80 (20 mg/kg, s.c.) | WT-Control: 15<br>WT-SNC80: 15<br>B2-Control: 21<br>B2-SNC80: 21 | Two-Way ANOVA test | Interaction<br>Genotype factor<br>Drug factor | F(1,68) = 0.3392<br>F(1,68) = 20.7<br>F(1.68) = 0.796 | 0.5622<br><0.0001<br>0.3754 | Sidak's Multiple Comparison Test | WT: Con vs. SNC80 | 7.9330 | 0.5615 | ns | |
| | | | | | | | | | | B2 KO: Con vs. SNC80 | 1.6670 | 0.9643 | ns | |
| Figure 1-g | Dark-light box test (Total transition) | WT & βarr2 KO | SNC80 (20 mg/kg, s.c.) | WT-Control: 12<br>WT-SNC80: 11<br>B2-Control: 20<br>B2-SNC80: 18 | Two-Way ANOVA test | Interaction<br>Genotype factor<br>Drug factor | F(1,57) = 1.754<br>F(1,57) = 0.1222<br>F(1.57) = 0.03687 | 0.1907<br>0.728<br>0.8484 | Sidak's Multiple Comparison Test | WT: Con vs. SNC80 | -2.6140 | 0.7254 | ns | |
| | | | | | | | | | | B2 KO: Con vs. SNC80 | 3.5000 | 0.3949 | ns | |
| Figure S1-a | Dark-light box test (Total distance -cm) | WT & βarr2 KO | SNC80 (20 mg/kg, s.c.) | WT-Control: 12<br>WT-SNC80: 11<br>B2-Control: 20<br>B2-SNC80: 20 | Two-Way ANOVA test | Interaction<br>Genotype factor<br>Drug factor | F (1, 59) = 1.949<br>F (1, 59) = 15.40<br>F (1, 59) = 19.78 | P=0.1679<br>P=0.0002<br>P<0.0001 | Sidak's Multiple Comparison Test | WT: Con vs. SNC80 | -390.8000 | 0.0011 | ** | |
| | | | | | | | | | | B2 KO: Con vs. SNC80 | -204.1000 | 0.0283 | * | |

| Subfigure | Brain region | Genotype | Drug | # of samples | Test | F-value | p-value | Group | Mean | Post hoc analysis | Group Comparison | Mean Diff. | p-value | Significance |
|---|---|---|---|---|---|---|---|---|---|---|---|---|---|---|
| Figure S1-b | Dar-light box test (Total distance - cm) | WT | SNC80 (20 mg/kg, i.p) or SL327 (50 mg/kg s.c.) | Con: 19<br>SNC80: 10<br>SNC+SL: 12<br>SL327: 12 | One-Way ANOVA test | F (3, 49) = 9.037 | P<0.0001 | Control<br>SNC80<br>SNC+SL327<br>SL327 | 811.8<br>1090<br>663.2<br>705.8 | Tukey's Multiple Comparison Test | Control vs. SNC80<br>Control vs. SNC+SL327<br>Control vs. SL327<br>SNC80 vs. SNC+SL327<br>SNC80 vs. SL327<br>SNC+SL327 vs. SL327 | -278.1000<br>148.6000<br>106.1000<br>426.7000<br>384.2000<br>-42.5000 | 0.0067<br>0.2258<br>0.5155<br><0.0001<br>0.0004<br>0.9585 | **<br>ns<br>ns<br>****<br>***<br>ns |

**Table S2. Statistical analysis of ERK1/2 expression levels upon time-series administration of SNC80 in WT and β-arrestin 2 KO mouse brain** Statistical differences of ERK1/2 expression levels in WT mice shown in **Fig. 2** and β-arrestin 2 KO mice in **Fig. 3**. Significance between groups was calculated by one-way ANOVA followed by a Tukey's multiple comparison (*$p$<0.05, **$p$<0.01, and ns=not significant).

| Subfigure | Brain region | Genotype | Drug | # of samples | Test | F-value | p-value | Group | Mean | Post hoc analysis | Group Comparison | Mean Diff. | p-value | Significance |
|---|---|---|---|---|---|---|---|---|---|---|---|---|---|---|
| Figure 2-e | Dorsal Striatum | WT | SNC80 (20 mg/kg, i.p) | Con: 13<br>10 min: 13<br>30 min: 5 | One-Way ANOVA test | F (2,28) = 6.776 | P=0.0040 | Con | 1 | Tukey's Multiple Comparison Test | Con vs. 10 min | -0.4718 | 0.0128 | * |
| | | | | | | | | 10 min | 1.472 | | Con vs. 30 min | 0.1534 | 0.7398 | ns |
| | | | | | | | | 30 min | 0.8466 | | 10 min vs. 30 min | 0.6253 | 0.0140 | * |
| Figure 2-f | Nucleus Acumbens | WT | SNC80 (20 mg/kg, i.p) | Con: 13<br>10 min: 13<br>30 min: 6 | One-Way ANOVA test | F (2,29) = 6.645 | P=0.0042 | Con | 1 | Tukey's Multiple Comparison Test | Con vs. 10 min | -0.3632 | 0.0317 | * |
| | | | | | | | | 10 min | 1.363 | | Con vs. 30 min | 0.2070 | 0.4563 | ns |
| | | | | | | | | 30 min | 0.793 | | 10 min vs. 30 min | 0.5702 | 0.0064 | ** |
| Figure 2-g | Dorsal Hippocampus | WT | SNC80 (20 mg/kg, i.p) | Con: 13<br>10 min: 12<br>30 min: 6 | One-Way ANOVA test | F (2,28) = 8.252 | P=0.0015 | Con | 1 | Tukey's Multiple Comparison Test | Con vs. 10 min | -0.8194 | 0.0051 | ** |
| | | | | | | | | 10 min | 1.819 | | Con vs. 30 min | 0.1918 | 0.7923 | ns |
| | | | | | | | | 30 min | 0.8082 | | 10 min vs. 30 min | 1.0110 | 0.0056 | ** |
| Figure 2-h | Amygdala | WT | SNC80 (20 mg/kg, i.p) | Con: 13<br>10 min: 13<br>30 min: 6 | One-Way ANOVA test | F (2,29) = 10.82 | P=0.0003 | Con | 1 | Tukey's Multiple Comparison Test | Con vs. 10 min | -0.5264 | 0.0024 | ** |
| | | | | | | | | 10 min | 1.526 | | Con vs. 30 min | 0.1940 | 0.5284 | ns |
| | | | | | | | | 30 min | 0.806 | | 10 min vs. 30 min | 0.7204 | 0.0010 | ** |
| Figure 2-i | Ventral Hippocampus | WT | SNC80 (20 mg/kg, i.p) | Con: 11<br>10 min: 10<br>30 min: 3 | One-Way ANOVA test | F (2,21) = 2.82 | P=0.0823 | Con | 1 | Tukey's Multiple Comparison Test | Con vs. 10 min | -0.1379 | 0.6248 | ns |
| | | | | | | | | 10 min | 1.138 | | Con vs. 30 min | 0.3892 | 0.2039 | ns |
| | | | | | | | | 30 min | 0.6108 | | 10 min vs. 30 min | 0.5271 | 0.0676 | ns |
| Figure 3-c | Dorsal Striatum | βarr2 KO | SNC80 (20 mg/kg, i.p) | Con: 8<br>10 min: 7<br>30 min: 8 | One-Way ANOVA test | F (2, 20) = 1.873 | P=0.1220 | Con | 1 | Tukey's Multiple Comparison Test | Con vs. 10 min | -0.7647 | 0.1805 | ns |
| | | | | | | | | 10 min | 1.765 | | Con vs. 30 min | -0.1857 | 0.8887 | ns |
| | | | | | | | | 30 min | 1.186 | | 10 min vs. 30 min | 0.5790 | 0.3608 | ns |
| Figure 3-d | Nucleus Acumbens | βarr2 KO | SNC80 (20 mg/kg, i.p) | Con: 7<br>10 min: 7<br>30 min: 7 | One-Way ANOVA test | F (2, 18) = 4.903 | P=0.0200 | Con | 1 | Tukey's Multiple Comparison Test | Con vs. 10 min | -0.8913 | 0.0240 | * |
| | | | | | | | | 10 min | 1.891 | | Con vs. 30 min | -0.1407 | 0.8907 | ns |
| | | | | | | | | 30 min | 1.141 | | 10 min vs. 30 min | 0.7506 | 0.0608 | ns |
| Figure 3-e | Dorsal Hippocampus | βarr2 KO | SNC80 (20 mg/kg, i.p) | Con: 8<br>10 min: 7<br>30 min: 8 | One-Way ANOVA test | F (2, 20) = 0.8178 | P=0.4556 | Con | 1 | Tukey's Multiple Comparison Test | Con vs. 10 min | -0.1654 | 0.5183 | ns |
| | | | | | | | | 10 min | 1.165 | | Con vs. 30 min | 0.0024 | 0.9998 | ns |
| | | | | | | | | 30 min | 0.9976 | | 10 min vs. 30 min | 0.1678 | 0.5086 | ns |
| Figure 3-f | Amygdala | βarr2 KO | SNC80 (20 mg/kg, i.p) | Con: 8<br>10 min: 8<br>30 min: 8 | One-Way ANOVA test | F (2, 21) = 1.472 | P=0.2522 | Con | 1 | Tukey's Multiple Comparison Test | Con vs. 10 min | -0.2360 | 0.4909 | ns |
| | | | | | | | | 10 min | 1.236 | | Con vs. 30 min | 0.1058 | 0.8631 | ns |
| | | | | | | | | 30 min | 0.8942 | | 10 min vs. 30 min | 0.3418 | 0.2376 | ns |
| Figure 3-g | Ventral Hippocampus | βarr2 KO | SNC80 (20 mg/kg, i.p) | Con: 8<br>10 min: 7<br>30 min: 8 | One-Way ANOVA test | F (2, 20) = 2.035 | P=0.1569 | Con | 1 | Tukey's Multiple Comparison Test | Con vs. 10 min | -0.2905 | 0.2167 | ns |
| | | | | | | | | 10 min | 1.291 | | Con vs. 30 min | 0.0098 | 0.9980 | ns |
| | | | | | | | | 30 min | 0.9902 | | 10 min vs. 30 min | 0.3003 | 0.1968 | ns |

**Table S3. Statistical analysis of anxiety-like behavior and ERK1/2 expression levels upon administration of SNC80 in the presence/absence of SL327 in WT mouse brain** Statistical differences of anxiety-like behavior and ERK1/2 expression levels in WT mouse brain shown in **Fig. 4** and **Fig. S3**. Significance between groups was calculated by one-way ANOVA followed by a Tukey's multiple comparison (**p*<0.05, ***p*<0.01, ****p*<0.001, and ns=not significant).

| Subfigure | Behavior test or Brain region | Genotype | Drug | # of samples | Test | F-value | p-value | Group | Mean | Post hoc analysis | Group Comparison | Mean Diff. | p-value | Significance |
|---|---|---|---|---|---|---|---|---|---|---|---|---|---|---|
| Figure 4-b | Dar-light box test | WT | SNC80 (20 mg/kg, i.p) or SL327 (50 mg/kg s.c.) | Con: 7<br>SNC80: 9<br>SNC+SL: 11<br>SL327: 11 | One-Way ANOVA test | F (3, 34) = 12.35 | P<0.0001 | Control<br>SNC80<br>SNC+SL327<br>SL327 | 13.88<br>30.53<br>3.418<br>15.64 | Tukey's Multiple Comparison Test | Control vs. SNC80 | -16.6400 | 0.0109 | * |
| | | | | | | | | | | | Control vs. SNC+SL327 | 10.4600 | 0.1496 | ns |
| | | | | | | | | | | | Control vs. SL327 | -1.7550 | 0.9830 | ns |
| | | | | | | | | | | | SNC80 vs. SNC+SL327 | 27.1100 | <0.0001 | **** |
| | | | | | | | | | | | SNC80 vs. SL327 | 14.8900 | 0.0106 | * |
| | | | | | | | | | | | SNC+SL327 vs. SL327 | -12.2200 | 0.0325 | * |
| Figure 4-c | Dorsal Hippocampus | WT | SNC80 (20 mg/kg, i.p) or SL327 (50 mg/kg s.c.) | Con: 10<br>SNC80: 9<br>SNC+SL: 9<br>SL327: 9 | One-Way ANOVA test | F (3, 33) = 11.68 | P<0.0001 | Control<br>SNC80<br>SNC+SL327<br>SL327 | 1<br>1.706<br>0.9214<br>0.7122 | Tukey's Multiple Comparison Test | Control vs. SNC80 | -0.6767 | 0.0024 | ** |
| | | | | | | | | | | | Control vs. SNC+SL327 | 0.1076 | 0.9247 | ns |
| | | | | | | | | | | | Control vs. SL327 | 0.3169 | 0.2790 | ns |
| | | | | | | | | | | | SNC80 vs. SNC+SL327 | 0.7843 | 0.0006 | *** |
| | | | | | | | | | | | SNC80 vs. SL327 | 0.9936 | <0.0001 | **** |
| | | | | | | | | | | | SNC+SL327 vs. SL327 | 0.2092 | 0.6459 | ns |
| Figure 4-d | Amygdala | WT | SNC80 (20 mg/kg, i.p) or SL327 (50 mg/kg s.c.) | Con: 10<br>SNC80: 10<br>SNC+SL: 10<br>SL327: 9 | One-Way ANOVA test | F (3, 35) = 10.9 | P<0.0001 | Control<br>SNC80<br>SNC+SL327<br>SL327 | 1<br>2.003<br>1.133<br>0.5908 | Tukey's Multiple Comparison Test | Control vs. SNC80 | -0.9798 | 0.0021 | ** |
| | | | | | | | | | | | Control vs. SNC+SL327 | -0.1093 | 0.9713 | ns |
| | | | | | | | | | | | Control vs. SL327 | 0.4325 | 0.3442 | ns |
| | | | | | | | | | | | SNC80 vs. SNC+SL327 | 0.8705 | 0.0069 | ** |
| | | | | | | | | | | | SNC80 vs. SL327 | 1.4120 | <0.0001 | **** |
| | | | | | | | | | | | SNC+SL327 vs. SL327 | 0.5419 | 0.1677 | ns |
| Figure S3-a | Dorsal Striatum | WT | SNC80 (20 mg/kg, i.p) or SL327 (50 mg/kg s.c.) | Con: 9<br>SNC80: 9<br>SNC+SL: 9<br>SL327: 9 | One-Way ANOVA test | F (3, 32) = 6.421 | P=0.0016 | Control<br>SNC80<br>SNC+SL327<br>SL327 | 1<br>1.543<br>1.115<br>0.7734 | Tukey's Multiple Comparison Test | Control vs. SNC80 | -0.5429 | 0.0247 | * |
| | | | | | | | | | | | Control vs. SNC+SL327 | -0.1155 | 0.9180 | ns |
| | | | | | | | | | | | Control vs. SL327 | 0.2266 | 0.5960 | ns |
| | | | | | | | | | | | SNC80 vs. SNC+SL327 | 0.4274 | 0.1032 | ns |
| | | | | | | | | | | | SNC80 vs. SL327 | 0.7694 | 0.0009 | *** |
| | | | | | | | | | | | SNC+SL327 vs. SL327 | 0.3420 | 0.2490 | ns |
| Figure S3-b | Nucleus Accumbens | WT | SNC80 (20 mg/kg, i.p) or SL327 (50 mg/kg s.c.) | Con: 10<br>SNC80: 10<br>SNC+SL: 10<br>SL327: 9 | One-Way ANOVA test | F (3, 35) = 2.672 | P=0.0624 | Control<br>SNC80<br>SNC+SL327<br>SL327 | 1<br>1.319<br>1.04<br>0.7877 | Tukey's Multiple Comparison Test | Control vs. SNC80 | -0.3193 | 0.3248 | ns |
| | | | | | | | | | | | Control vs. SNC+SL327 | -0.0404 | 0.9963 | ns |
| | | | | | | | | | | | Control vs. SL327 | 0.2123 | 0.6809 | ns |
| | | | | | | | | | | | SNC80 vs. SNC+SL327 | 0.2790 | 0.4427 | ns |
| | | | | | | | | | | | SNC80 vs. SL327 | 0.5316 | 0.0394 | * |
| | | | | | | | | | | | SNC+SL327 vs. SL327 | 0.2526 | 0.5501 | ns |
| Figure S3-c | Ventral Hippocampus | WT | SNC80 (20 mg/kg, i.p) or SL327 (50 mg/kg s.c.) | Con: 10<br>SNC80: 9<br>SNC+SL: 10<br>SL327: 9 | One-Way ANOVA test | F (3, 34) = 2.734 | P=0.0588 | Control<br>SNC80<br>SNC+SL327<br>SL327 | 1<br>1.164<br>0.7771<br>0.8315 | Tukey's Multiple Comparison Test | Control vs. SNC80 | -0.1522 | 0.7419 | ns |
| | | | | | | | | | | | Control vs. SNC+SL327 | 0.2347 | 0.3879 | ns |
| | | | | | | | | | | | Control vs. SL327 | 0.1803 | 0.6299 | ns |
| | | | | | | | | | | | SNC80 vs. SNC+SL327 | 0.3869 | 0.0655 | ns |
| | | | | | | | | | | | SNC80 vs. SL327 | 0.3325 | 0.1549 | ns |
| | | | | | | | | | | | SNC+SL327 vs. SL327 | -0.0545 | 0.9833 | ns |

**Table S4. Statistical analysis of fear-related behavior upon systemic administration of SNC80 or TAN67 in WT and β-arrestin 2 KO mice** Statistical differences of fear-related behaviors in WT or β-arrestin 2 KO mice shown in Fig. 5. Significance between groups was calculated by two-way ANOVA followed by a Bonferroni's multiple comparison (**p*<0.05, *****p*<0.0001, and ns=not significant).

| Subfigure | Behavior test | Genotype | Drug | # of samples | Test | Source of Variation | F-value | p-value | Post hoc analysis | Group Comparison | Mean Diff. | p-value | Significance |
|---|---|---|---|---|---|---|---|---|---|---|---|---|---|
| Figure 5-c | Fear potentiated startle test (Raw startle) | WT | SNC80 (20 mg/kg, i.p.) | Control: 21<br>SNC80: 21 | Two-Way ANOVA test | Interaction | F(2,120) = 20.42 | <0.0001 | Sidak's Multiple Comparison Test | Blank: Con vs. SNC80 | -0.0126 | 0.9994 | ns |
| | | | | | | Stimulation factor | F(2,120) = 92.80 | <0.0001 | | Noise: Con vs. SNC80 | 0.5879 | <0.0001 | **** |
| | | | | | | Drug factor | F(1,120) = 63.99 | <0.0001 | | Noise+Light: Con vs. SNC80 | 1.0310 | <0.0001 | **** |
| Figure 5-d | Fear potentiated startle test (FPS testing) | WT | SNC80 (20 mg/kg, i.p.) | Control: 21<br>SNC80: 20 | Unpaird t test | | | p=0.0085 | | | | | |
| Figure 5-f | Fear potentiated startle test | βarr2 KO | SNC80 (20 mg/kg, i.p.) | Control: 8<br>SNC80: 8 | Two-Way ANOVA test | Interaction | F(2,42) = 20.22 | <0.0001 | Sidak's Multiple Comparison Test | Blank: Con vs. SNC80 | -0.0355 | >0.9999 | ns |
| | | | | | | Stimulation factor | F(2,42) = 51.52 | <0.0001 | | Noise: Con vs. SNC80 | 0.2139 | 0.0103 | * |
| | | | | | | Drug factor | F(1,42) = 40.4 | <0.0001 | | Noise+Light: Con vs. SNC80 | 0.5812 | <0.0001 | **** |
| Figure 5-g | Fear potentiated startle test (FPS testing) | βarr2 KO | SNC80 (20 mg/kg, i.p.) | Control: 8<br>SNC80: 8 | Unpaird t test | | | p=0.0003 | | | | | |
| Figure 5-i | Fear potentiated startle test | WT | TAN67 (25 mg/kg, i.p.) | Control:8<br>TAN67: 8 | Two-Way ANOVA test | Interaction | F(2,42) = 0.7245 | 0.4905 | Sidak's Multiple Comparison Test | Blank: Con vs. SNC80 | -0.0113 | >0.9999 | ns |
| | | | | | | Stimulation factor | F(2,42) = 25.06 | <0.0001 | | Noise: Con vs. SNC80 | 0.1123 | 0.9736 | ns |
| | | | | | | Drug factor | F(1,42) = 0.2754 | 0.6025 | | Noise+Light: Con vs. SNC80 | -0.3664 | 0.5189 | ns |
| Figure 5-j | Fear potentiated startle test (FPS testing) | WT | TAN67 (25 mg/kg, i.p.) | Control: 7<br>TAN67: 8 | Unpaird t test | | | p<0.0001 | | | | | |

**Table S5. Statistical analysis of ERK1/2 expression levels upon time-series administration of TAN67 in WT mousee and SNC80 in β-arrestin 1 KO mouse brain** Statistical differences of ERK1/2 expression levels in WT and β-arrestin 1 KO mouse brain shown in **Fig. 6**. Significance between groups was calculated by one-way ANOVA followed by a Tukey's multiple comparison (*$p$<0.05, **$p$<0.01, and ns=not significant).

| Subfigure | Brain region | Genotype | Drug | # of samples | Test | F-value | p-value | Group | Mean | Post hoc analysis | Group Comparison | Mean Diff. | p-value | Significance |
|---|---|---|---|---|---|---|---|---|---|---|---|---|---|---|
| Figure 6-a | Dorsal Striatum | WT | TAN67 (25 mg/kg, i.p) | Con: 7<br>10 min: 7<br>30 min: 7 | One-Way ANOVA test | F (2, 18) = 5.276 | P=0.0157 | Con<br>10 min<br>30 min | 1<br>0.8769<br>0.5824 | Tukey's Multiple Comparison Test | Con vs. 10 min<br>Con vs. 30 min<br>10 min vs. 30 min | 0.1231<br>0.4176<br>0.2945 | 0.6279<br>0.0142<br>0.0931 | ns<br>*<br>ns |
| Figure 6-b | Nucleus Acumbens | WT | TAN67 (25 mg/kg, i.p) | Con: 7<br>10 min: 7<br>30 min: 6 | One-Way ANOVA test | F (2, 17) = 5.701 | P=0.0127 | Con<br>10 min<br>30 min | 1<br>0.6152<br>0.5227 | Tukey's Multiple Comparison Test | Con vs. 10 min<br>Con vs. 30 min<br>10 min vs. 30 min | 0.3848<br>0.4773<br>0.0925 | 0.0447<br>0.0161<br>0.8190 | *<br>*<br>ns |
| Figure 6-c | Dorsal Hippocampus | WT | TAN67 (25 mg/kg, i.p) | Con: 7<br>10 min: 7<br>30 min: 7 | One-Way ANOVA test | F (2, 18) = 5.09 | P=0.0177 | Con<br>10 min<br>30 min | 1<br>1.021<br>0.739 | Tukey's Multiple Comparison Test | Con vs. 10 min<br>Con vs. 30 min<br>10 min vs. 30 min | -0.0207<br>0.2610<br>0.2817 | 0.9759<br>0.0410<br>0.0266 | ns<br>*<br>* |
| Figure 6-d | Amygdala | WT | TAN67 (25 mg/kg, i.p) | Con: 7<br>10 min: 6<br>30 min: 6 | One-Way ANOVA test | F (2, 16) = 5.455 | P=0.0156 | Con<br>10 min<br>30 min | 1<br>0.851<br>0.6697 | Tukey's Multiple Comparison Test | Con vs. 10 min<br>Con vs. 30 min<br>10 min vs. 30 min | 0.1490<br>0.3303<br>0.1813 | 0.3216<br>0.0118<br>0.2189 | ns<br>*<br>ns |
| Figure 6-e | Ventral Hippocampus | WT | TAN67 (25 mg/kg, i.p) | Con: 6<br>10 min: 6<br>30 min: 6 | One-Way ANOVA test | F (2, 15) = 1.092 | P=0.3607 | Con<br>10 min<br>30 min | 1<br>1.063<br>0.8009 | Tukey's Multiple Comparison Test | Con vs. 10 min<br>Con vs. 30 min<br>10 min vs. 30 min | -0.0628<br>0.1991<br>0.2618 | 0.9388<br>0.5426<br>0.3582 | ns<br>ns<br>ns |
| Figure 6-f | Dorsal Striatum | βarr1 KO | SNC80 (20 mg/kg, i.p) | Con: 7<br>10 min: 6<br>30 min: 6 | One-Way ANOVA test | F (2,16) = 1.62 | P=0.2288 | Con<br>10 min<br>30 min | 1<br>1.191<br>1.271 | Tukey's Multiple Comparison Test | Con vs. 10 min<br>Con vs. 30 min<br>10 min vs. 30 min | -0.1907<br>-0.2710<br>-0.0803 | 0.4580<br>0.2226<br>0.8743 | ns<br>ns<br>ns |
| Figure 6-g | Nucleus Acumbens | βarr1 KO | SNC80 (20 mg/kg, i.p) | Con: 7<br>10 min: 6<br>30 min: 6 | One-Way ANOVA test | F (2,16) = 0.5181 | P=0.6053 | Con<br>10 min<br>30 min | 1<br>1.179<br>1.232 | Tukey's Multiple Comparison Test | Con vs. 10 min<br>Con vs. 30 min<br>10 min vs. 30 min | -0.1791<br>-0.2319<br>-0.0528 | 0.7426<br>0.6112<br>0.9759 | ns<br>ns<br>ns |
| Figure 6-h | Dorsal Hippocampus | βarr1 KO | SNC80 (20 mg/kg, i.p) | Con: 7<br>10 min: 6<br>30 min: 6 | One-Way ANOVA test | F (2,16) = 11.48 | P=0.0008 | Con<br>10 min<br>30 min | 1<br>2.812<br>2.874 | Tukey's Multiple Comparison Test | Con vs. 10 min<br>Con vs. 30 min<br>10 min vs. 30 min | -1.8120<br>-1.8740<br>-0.0625 | 0.0027<br>0.0020<br>0.9902 | **<br>**<br>ns |
| Figure 6-i | Amygdala | βarr1 KO | SNC80 (20 mg/kg, i.p) | Con: 7<br>10 min: 6<br>30 min: 6 | One-Way ANOVA test | F (2,16) = 4.668 | P=0.0253 | Con<br>10 min<br>30 min | 1<br>1.498<br>1.67 | Tukey's Multiple Comparison Test | Con vs. 10 min<br>Con vs. 30 min<br>10 min vs. 30 min | -0.4981<br>-0.6695<br>-0.1714 | 0.1080<br>0.0261<br>0.7564 | ns<br>*<br>ns |
| Figure 6-j | Ventral Hippocampus | βarr1 KO | SNC80 (20 mg/kg, i.p) | Con: 7<br>10 min: 6<br>30 min: 6 | One-Way ANOVA test | F (2,16) = 5.097 | P=0.0194 | Con<br>10 min<br>30 min | 1<br>2.024<br>2.377 | Tukey's Multiple Comparison Test | Con vs. 10 min<br>Con vs. 30 min<br>10 min vs. 30 min | -1.0240<br>-1.3770<br>-0.3530 | 0.0907<br>0.0201<br>0.7372 | ns<br>*<br>ns |

**Table S6. Antibody information for the Western blot** Lists of primary and secondary antibodies that were used in the study were included in the table.

| Name of primary antibody | Company | Molecular Weight (kDa) | Source | Dilution ratio | Catalog number | Lot number |
|---|---|---|---|---|---|---|
| p44/42 MAPK (Erk1/2) (L34F12) | Cell Signaling, MA | 42, 44 | Mouse | 1:2,000 for WB; 1:250 for IF | 4696S | 22 |
| phospho-ERK1/2 (Tyr 204) | Santa Cruz Biotechnology, Dallas, TX | 42, 44 | Rabbit | 1:2,000 for WB | 7976-R | C1113 |
| Phospho-p44/42 MAPK (Erk1/2) (Thr202/Tyr204) (D13.14.4E) XP® | Cell Signaling, MA | 42, 44 | Rabbit | 1:2,000 for WB; 1:200 for IF | 4370S | 24 |
| p38 MAPK (D13E1) XP® | Cell Signaling, MA | 38 | Rabbit | 1:2,000 | 8690S | 6 |
| Phospho-p38 MAPK (Thr180/Tyr182) | Cell Signaling, MA | 38 | Rabbit | 1:2,000 | 9211S | 23 |
| JNK (D-2) | Santa Cruz Biotechnology, Dallas, TX | 46, 54 | Mouse | 1:2,000 | 7345 | L3015 |
| p-JNK (G-7) | Santa Cruz Biotechnology, Dallas, TX | 46, 54 | Mouse | 1:2,000 | 6254 | B2117 |
| α-Tubulin | Santa Cruz Biotechnology, Dallas, TX | 50 | Mouse | 1:2,000 | 5286 | G3117 |
| **Name of secondary antibody** | **Company** | **Molecular Weight (kDa)** | **Source** | **Dilution ratio** | **Catalog number** | **Lot number** |
| IRDye® 680LT | Li-Cor, Lincoln, NE | - | Mouse | 1:5,000 | 926-68020 | 60824-02 |
| IRDye® 800CW | Li-Cor, Lincoln, NE | - | Rabbit | 1:5,000 | 926-32211 | C61103-06 |
| Alexa fluor 594 Goat Anti-Rabbit IgG (H+L) Antibody | Life Technologies (Thermo Fisher), Waltham, MA | - | Rabbit | 1:1,000 | A-11012 | - |
| Alexa Fluor 488 Goat Anti-Mouse IgG (H+L) Antibody | Life Technologies (Thermo Fisher), Waltham, MA | - | Mouse | 1:1,000 | A11001 | - |

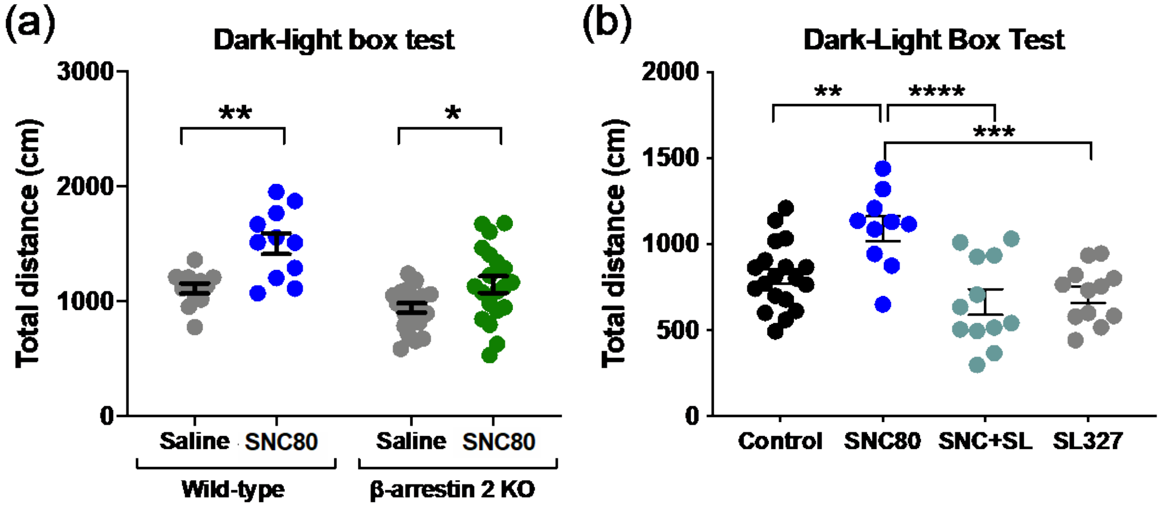


**Figure S1. Locomotor effects of drug/vehicle treatment in the WT or β-arrestin 2 KO mice in the dark light upon administration of drugs (a)** Traveled distance of WT (control: n=12, SNC80: n=11) and β-arrestin 2 KO mice (control: n=20, SNC80: n=20) upon administration of SNC80 (20 mg/kg, i.p.) the dark light box test shown in **Fig. 1e,g**. **(b)** Traveled distance of WT mice upon administration of SNC80 (20 mg/kg, i.p. / control: n=8, SNC80: n=12, SNC+SL: n=12, SL327: n=12) in presence or absence of 50 mg/kg SL327 in dark light box test shown in **Fig. 4b**. SNC80-induced hyperlocomotion corresponds with a previous report (*11*). (For (a), Significance was calculated by two-way ANOVA $F_{1,59}$=1.949, *p*=0.1670, WT *p*=0.0011, β-arrestin 2 KO *p*=0.0283 after Sidak's multiple comparison; for (b), one-way ANOVA $F_{3,49}$=9.037, *p*<0.001, control vs. SNC80 *p*=0.007, SNC80 vs. SNC+SL *p*<0.0001, SNC+SL vs. SL327 *p*<0.0004 followed by a Tukey's multiple comparison; **p*<0.05, ***p*<0.01, ****p*<0.001, *****p*<0.0001; all values are shown as individual data points ± S.E.M.).

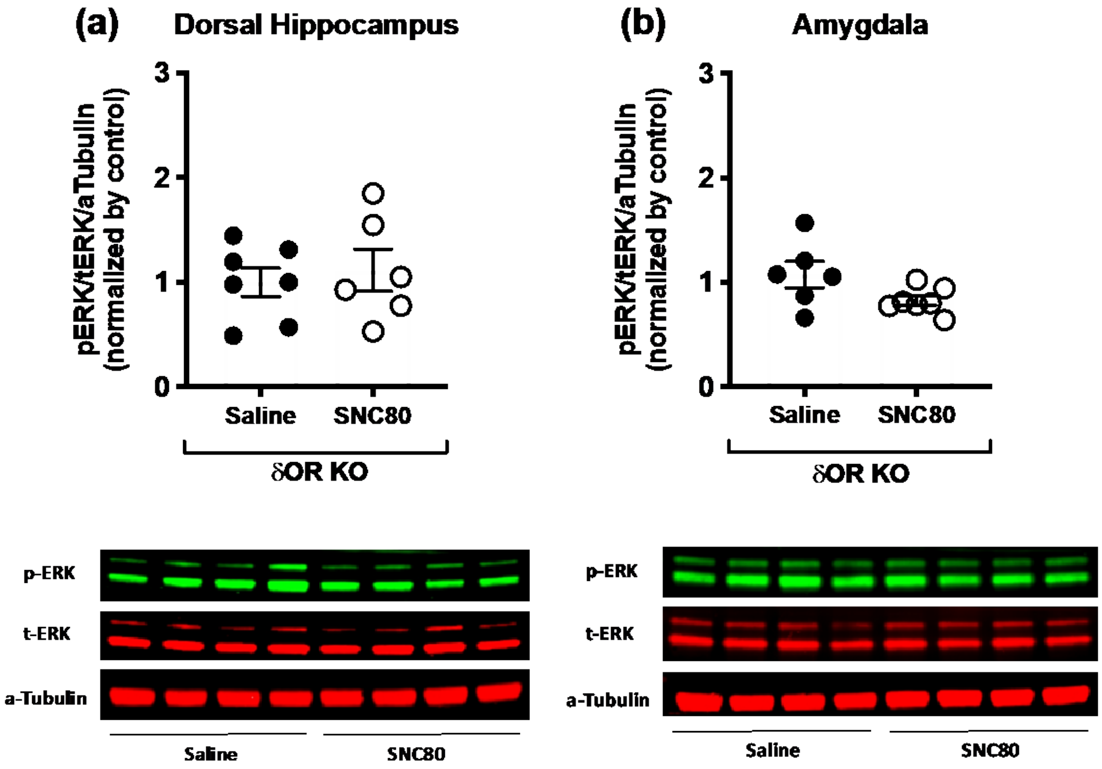


**Figure S2. SNC80 does not activate ERK1/2 in the dorsal hippocampus and the amygdala of δOR KO mice (a, b)** Unlike Fig. 2g,h, systemic administration of SNC80 (20 mg/kg, i.p.) 10 minutes prior to the brain tissue collections did not affect ERK1/2 activation profile in the dorsal hippocampus (Saline: n=8, SNC80: n=7) and the amygdala (Saline: n=7, SNC80: n=8) of δOR KO mice.

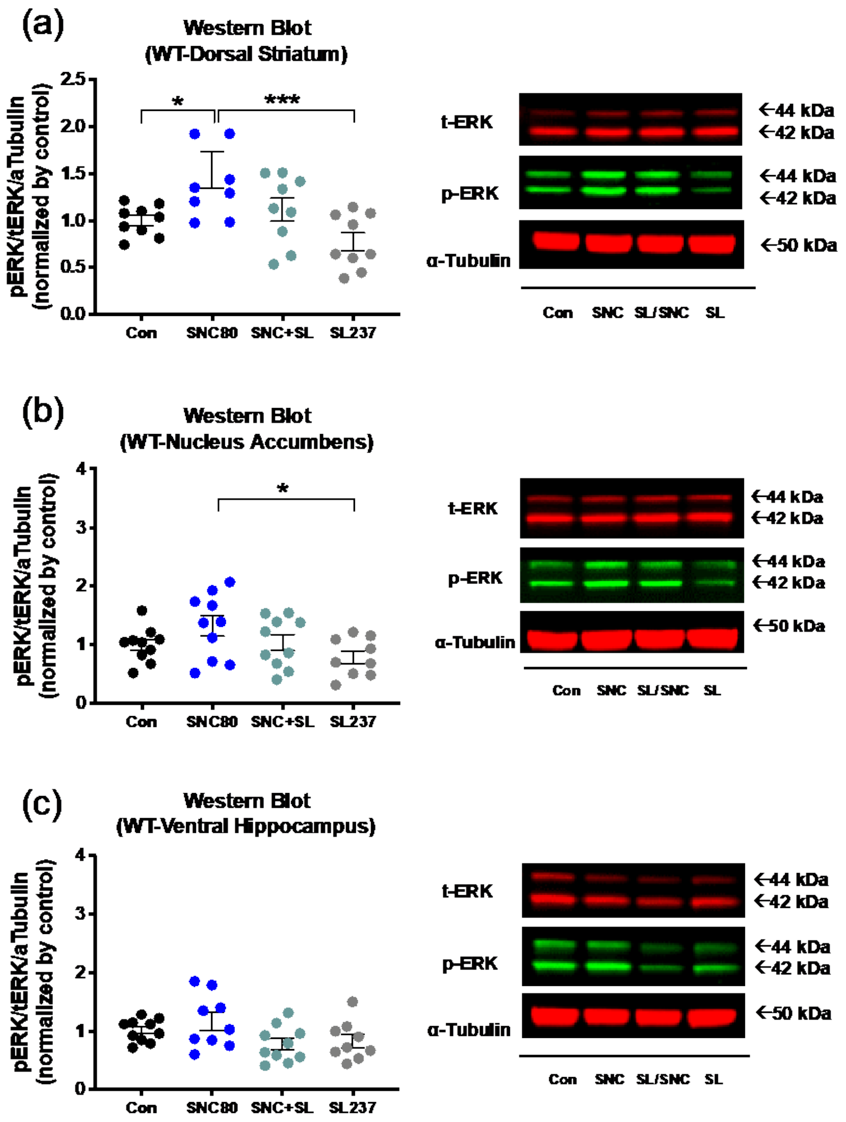


**Figure S3. SNC80-induced ERK1/2 activation is partly affected by SL327 in the striatal regions of the brain** SL327 (50 mg/kg, s.c.) attenuated SNC80 (20 mg/kg, i.p.)-induced ERK1/2 phosphorylation in the striatum similar to Fig. 4c,d **(a)** and similar trends were observed in the nucleus accumbens **(b)**. **(c)** Yet, no change was observed in the ventral hippocampus similar to Fig. 2i. The number of samples is listed in Table S3. (Significance was calculated by one-way ANOVA followed by a Sidak's or Tukey's multiple comparison; *$p$<0.05, ***$p$<0.001; all values are shown as individual data points ± S.E.M.; SNC+SL means SNC80+SL327 and SL means SL327).

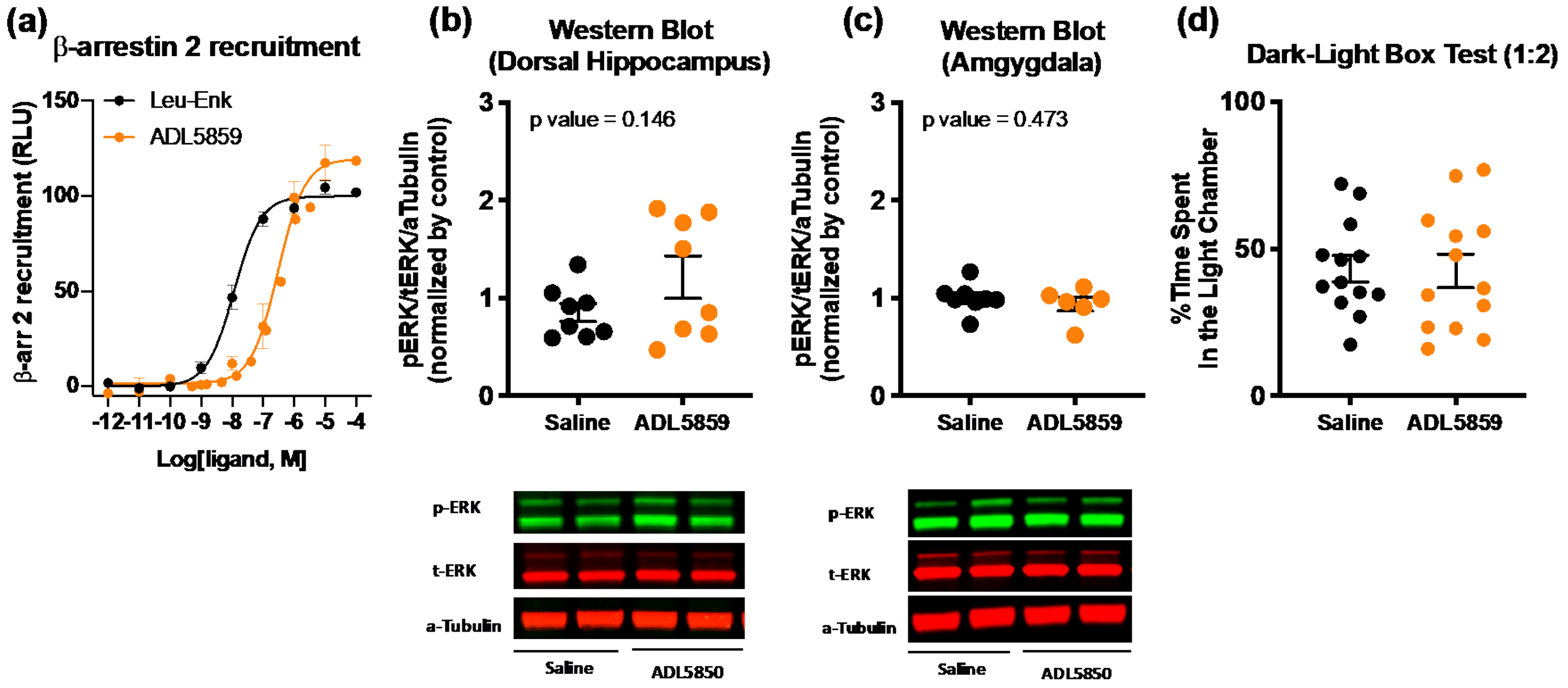


**Figure S4. A δOR agonist, ADL5859, does not affect ERK1/2 activity and anxiety-like behaviors of WT mice (a)** Dose-dependent β-arrestin 2 recruiment levels by ADL5859 and leucine-enkephalin (leu-enk) were evaluated using cellular assays in CHO-δOR-βarr2 cells (All recruiment levels were normalized by leu-enk and leu-enk was normalized as 100 %). **(b, c)** Systemic administration of ADL5859 (30 mg/kg, p.o.) did not affect ERK1/2 activation profile in the dorsal hippocampus (Saline: n=8, SNC80: n=8) and the amygdala (Saline: n=8, SNC80: n=6) of WT mice. ADL5859 was administered 10 minutes prior to the brain tissue collection. **(d)** No changes in anxiety-like behaviors in the dark/light box test were observed by systemic administration of ADL5859 (30 mg/kg, p.o.) (Saline: n=13, SNC80: n=13). ADL5859 was administered 30 mintues prior to the behavior testing.

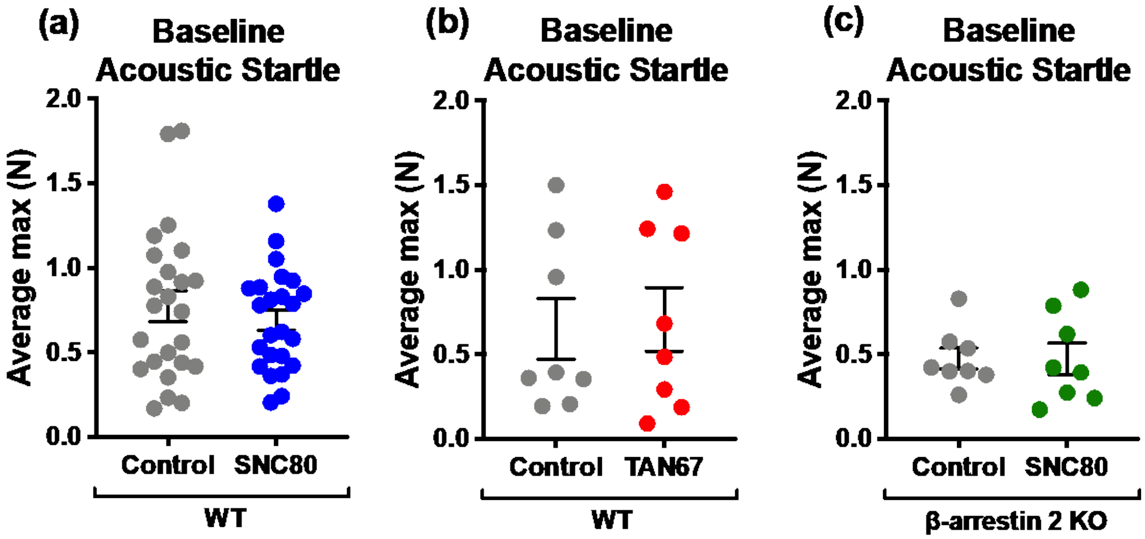


**Figure S5. Mice groups for FPS tests were counterbalanced based on baseline acoustic startle response (a)** No significance was observed between groups of control *vs.* SNC80 (Control: n=24, SNC80: n=24) **(b)** or control *vs.* TAN67 (Control: n=8, TAN67: n=8) of WT mice. **(b)** Also no significance was observed between control *vs.* SNC80 of β-arrestin 2 KO mice (Control: n=8, SNC80: n=8). All values are shown as individual data points ± S.E.M.

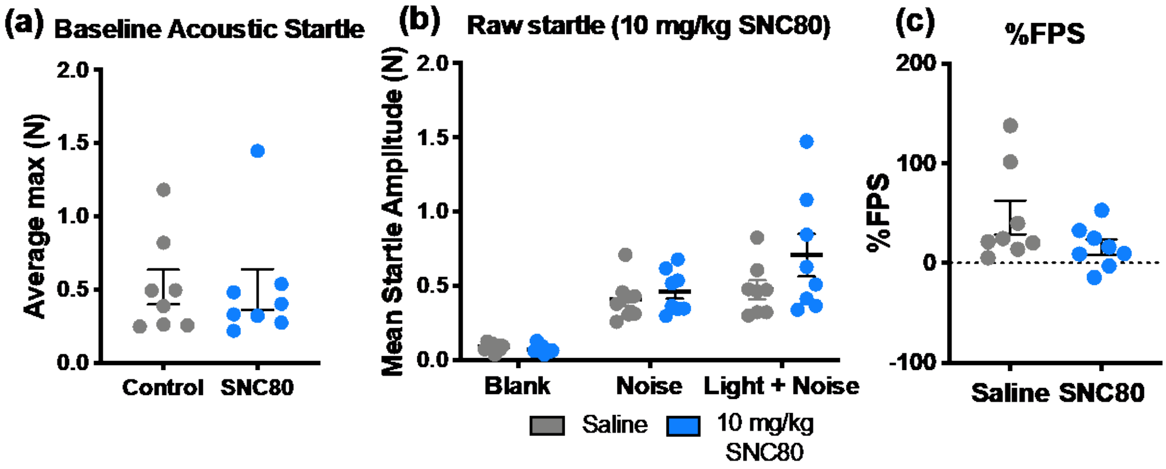


**Figure S6. A low dose SNC80 does not affect fear-related behavior of WT mice** Fear potentiated startle responses were evaluated upon administration of SNC80 (10 mg/kg, i.p.) in WT mice (Saline: n=8, SNC80: n=8). SNC80 was administered 30 mintues prior to the testing. **(a)** Prior to the testing, mice were measured with baseline acoustic startle and no difference was observed between groups (no drugs were administered for this period). **(b)** 10 mg/kg SNC80 did not affect the raw startle response to either 'noise' alone or 'light+noise' condition. **(c)** 10 mg/kg SNC80 also did not affect %FPS.

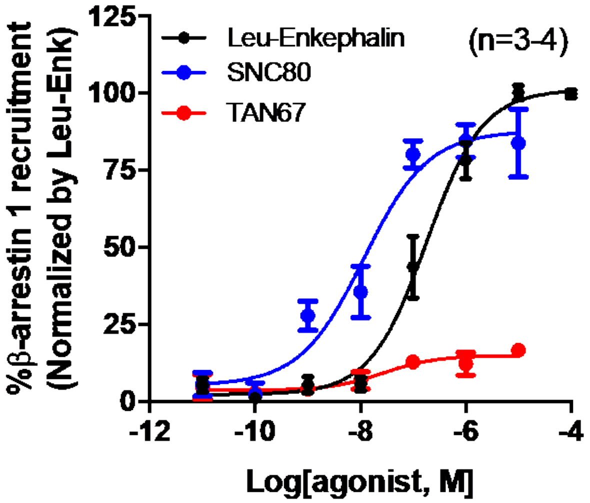

**Figure S7. β-arrestin 1 recruitment levels by G-protein-biased (TAN67), β-arrestin-biased (SNC80), and non-biased (Leu-Enk) δOR agonist in U2OS-δOR-βArr1 cells** Dose-dependent β-arrestin 1 recruiment levels by TAN67 (n=3), SNC80 (n=3), and Leu-enkephalin (n=4) were evaluated using the cellular assay in U2OS-δOR-βArr1 cells. SNC80 revealed the highest efficacy of recruiment and TAN67 showed the lowest. (All recruiment levels were normalized by leu-enk and leu-enk was normalized as 100 %).

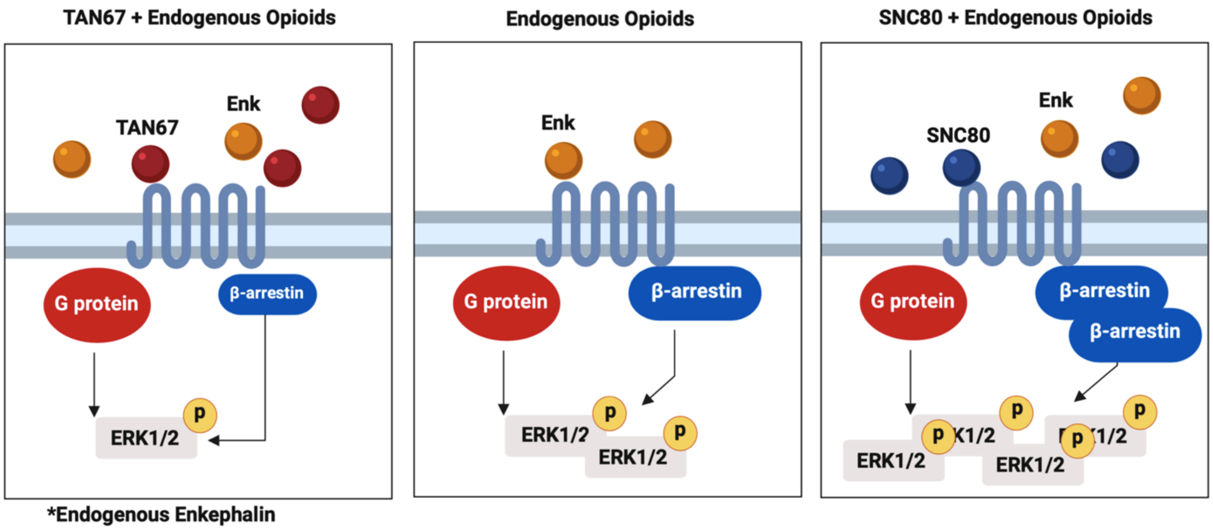


**Figure S8. A diagram respresenting the pharmacological competition between two biased agonists and an endnogenous opioid in relations to their ability to modulate ERK1/2 signaling** Unlike with cells, the brain has endogenous opioids that bind to δOR. As endogenous opioids such as Leu-Enk, an analog of endogenous opioids, have better ability to recruit β-arrestin proteins than TAN67 as shown in Fig. S7, δOR is less likely to recruit β-arrestin and potentially activate less ERK1/2 upon administration of TAN67 in the brain. Likewise, SNC80, which has a better ability to recruits β-arrestin proteins than Leu-Enk, recruits more β-arrestins via δOR and potentially activates more ERK1/2 upon administration of SNC80 in the brain (Right). Yet, it is notworthy that SNC80 and TAN67 have comparable levels of G protein-mediated response (*14*).
